# Structure of *D. melanogaster* ARC1 reveals a repurposed molecule with characteristics of retroviral Gag

**DOI:** 10.1101/691972

**Authors:** Matthew A. Cottee, Suzanne C. Letham, George R. Young, Jonathan P. Stoye, Ian A. Taylor

## Abstract

The tetrapod neuronal protein ARC and its *D. melanogaster* homologue, dARC1, have important but differing roles in neuronal development. Both are thought to originate through exaptation of ancient Ty3/Gypsy retrotransposon Gag genes, with their novel function relying on an original capacity for self-assembly and encapsidation of nucleic acids. Here, we present the crystal structure of dARC1 CA and examine the relationship between dARC1, mammalian ARC and the CA protein of circulating retroviruses. We show that whilst the overall architecture is highly related to that of orthoretroviral and spumaretroviral CA, there are significant deviations in both N- and C-terminal domains, potentially affecting recruitment of partner proteins and particle assembly. The degree of sequence and structural divergence suggests that Ty3/Gypsy Gag has been exapted on two separate occasions and that, although mammalian ARC and dARC1 share functional similarity, the structures have undergone different adaptations after appropriation into the tetrapod and insect genomes.

## INTRODUCTION

Activity-Regulated Cytoskeleton-Associated protein (ARC) is an immediate early gene product induced in response to high levels of synaptic activity and is directed to neuronal synapses through signalling sequences in its 3’ UTR (*1*). Mammalian ARC (mam-ARC) is essential for neuronal plasticity and is involved in memory (*2, 3*) acting as a regulator of AMPA receptors (*4, 5*). ARC has also been implicated in neurological disorders including Alzheimer’s disease (*6, 7*), Fragile-X syndrome (*8*) and schizophrenia (*9–11*). In *Drosophila melanogaster*, two homologues of mam-ARC are expressed, dARC1, and dARC2 (*12*). dARC1 is present at neuromuscular junctions and, along with its mRNA, has been implicated in regulating the behavioural starvation response but is not involved in synaptic plasticity (*13*). Therefore, comparing the structural and functional properties of mam-ARC and dARC1 might lead to a better understanding of cognition and memory consolidation.

The *ARC* gene is thought to be derived from the *gag* gene of a Ty3/Gypsy retrotransposon (*14*) that, subsequent to genomic insertion, has been repurposed to perform an advantageous function to the host (*15*). This connection between ARC and retrotransposons was made when sequence alignments revealed that the ARC proteins shared sequence similarity with the Gag protein of retroviruses or retrotransposons (*14*). These data also suggested that ARC is evolutionarily related to the Ty3/Gypsy family of retrotransposons. Further evidence came from crystal structures of two α-helical domains from *Rattus norvegicus* ARC (rARC) (*16*), which revealed that rARC N- and C-terminal capsid (CA) domains were structurally homologous to the N- and C-terminal CA domains of both *Orthoretrovirinae* (*16*) and also *Spumaretrovirinae* (*17*). Further phylogenetic analysis revealed that despite mam-ARC and dARC1 seemingly providing related functions in the host, dARC1 and the tetrapod ARCs most likely arose from separate lineages of Ty3/Gypsy, since dARC1 clustered with insect Ty3/Gypsy retrotransposons and tetrapod ARCs clustered with fish Ty3/Gypsy retrotransposons (*15*).

The relevance of ARC’s retrotransposon origin to its function in synaptic plasticity was not immediately obvious until the recent observation that mam-ARC and dARC1 can self-assemble into particles and package RNA for potential transfer between cells (*12, 15*), similarly to retrotransposons and retroviruses (*18, 19*). In *D. melanogaster*, it is proposed that dARC1 expressed at neuromuscular junction presynaptic boutons assembles into particles that encapsidate *dARC1* mRNA. Loaded particles might then be packaged and released as extracellular vesicles for intercellular transfer to the post synapse where mRNA release and translation can take place (*12, 15*), Similarly, mam-ARC can also encapsidate ARC mRNA into particles, allowing transfer from donor to recipient neurons, where ARC mRNA can be translated (*15*).

Since both dARC1 and mam-ARC are able to form capsid-like particles (*12, 15*), it seems likely that they share a degree of structurally similarity. To date, crystal structures of the individual domains from rARC have been determined (*16*) along with the solution NMR structure of the rARC CA (*20*). Here we report two crystal structures of the entire CA region of dARC1 at 1.7 Å and 2.3 Å and consider these structures in comparison to those of rARC and retroviral CA. dARC1 comprises two α-helical domains with a fold related to that observed in the CA-NtD and CA-CtD of orthoretroviral and spumaretroviral CA. However, we observe significant divergence in the NtD of dARC1 where an extended hydrophobic strand that packs against α1 and α3 of the core fold replaces the N-terminal β-hairpin and helix α1 found in orthoretroviral CAs. In the rARC structure, this hydrophobic strand is replaced by peptides from the binding partners Ca^2+^/calmodulin-dependent protein kinase 2A (CamK2A) and transmembrane AMPAR regulatory protein γ2 (TARPγ2) and may represent a functional adaptation for the recruitment of partner proteins. We also show that dARC1 utilises the same CtD-CtD interface required for assembly of retroviral CA into mature particles and propose that this obligate dimer represents a building block for dARC1 particle assembly. Further examination of the relationship between dARC1, mam-ARC and Gag from Ty retrotransposon families, reveals that, although dARC1 and mam-ARC are functional orthologues, the structural divergence in dARC1 and mam-ARC CA domains is consistent with the notion of Ty3/Gypsy Gag exaptation on two separate occasions. We suggest they may have undergone different adaptations after appropriation into the tetrapod and insect genomes.

## RESULTS

### Structures of dARC1 CA

We determined the crystal structure of the capsid domain region of dARC1, residues S39-N205 (dARC1 CA) using single wavelength anomalous diffraction (SAD) and crystals of Se-Met substituted protein. The structure was determined in both an orthorhombic and a hexagonal crystal form. The orthorhombic crystals diffracted to higher resolution allowing the structure to be refined to a final resolution of 1.7 Å with R-factor and Free R-factor of 18.1% and 21.3% respectively. Details of data collection, phasing and refinement are presented in **Table S1**. The asymmetric unit (ASU) contains two chains, each containing an α-helical N- (CA-NtD) and C-terminal domain (CA-CtD) (**Fig. 1A**). The chains are arranged in a dimer with a distinct U-shape reminiscent of a glacial trough (**Fig. 1A, right**). The CA-CtDs form the base of the trough and pack together to form a homodimer interface and the CA-NtDs form the sides of the trough and are separated by ∼45 Å. Inspection of each domain reveals the CA-NtD is made up from an extended N-terminal strand and a four-helix core (α1-α4) and the CA-CtD comprises a further five α-helix bundle (α5-α9) (**Fig. 1B, i, ii**). The tertiary folds of each domain are particularly similar and can be superimposed with an RMSD of 2.2 Å over 49 Cα atoms (**Fig. 1C)**. Moreover, it can be seen that the dARC1 N-terminal β-strand is topologically equivalent to α5 in the CtD, while NtD α1-4 are equivalent to CtD α6-9. This strong similarity of dARC1 CA domains provides further evidence for the notion that tandem domains of CA arose as the result of a gene duplication event (*17*). The hexagonal crystal form was independently solved and refined to a resolution of 2.3 Å and reveals an almost identical dimeric ASU that aligns with a RMSD of only 0.247 Å over 133 Cα pairs (**Fig. S1A to C**). Both structures appear especially stable around the CTD-mediated dimeric interface and when aligned through their CTDs show only small differences in the positioning of NtDs with respect to the CtDs (**Fig. S1D**).

**Fig. 1.**
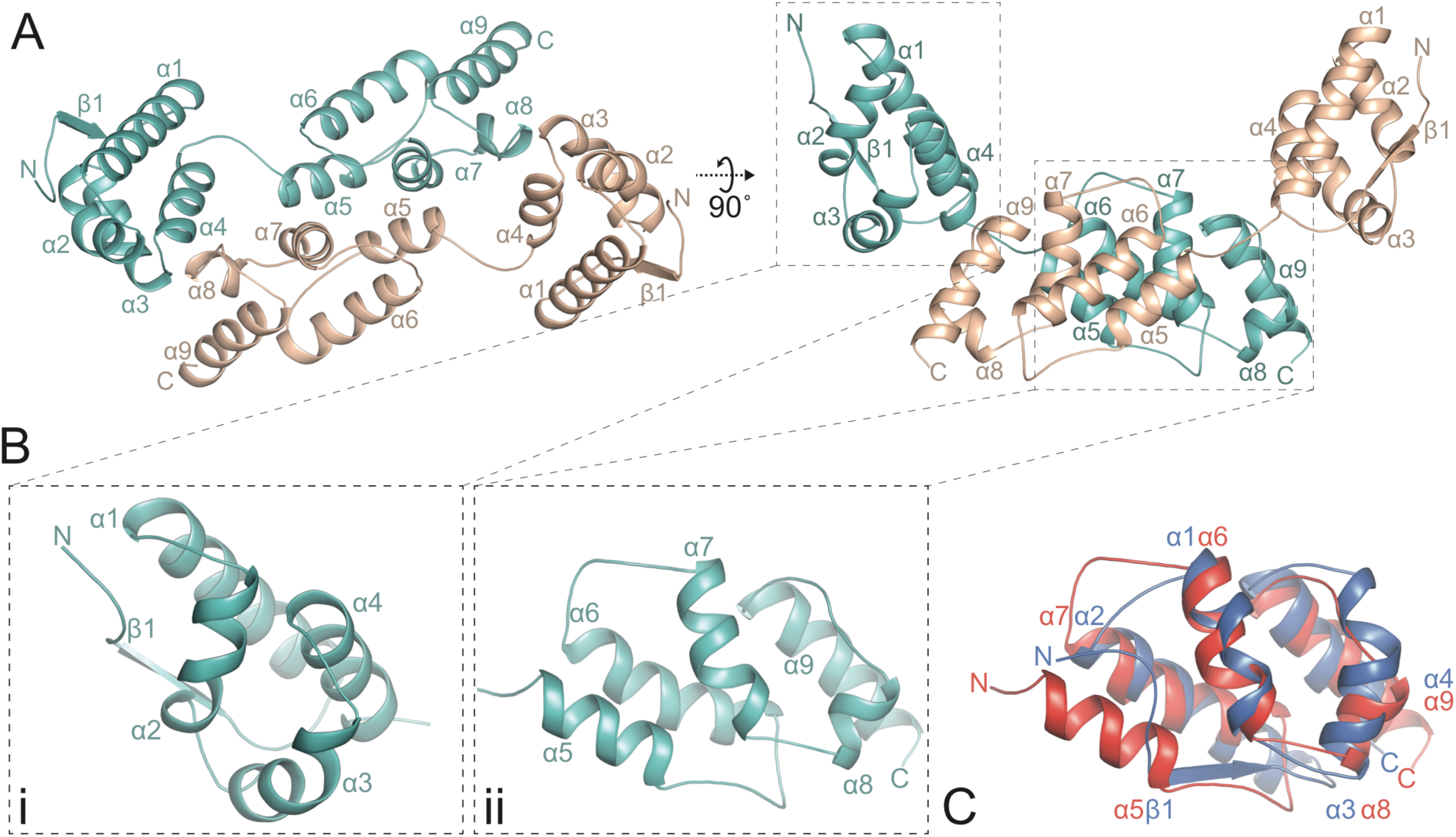
Crystal structure of the dARC1 CA domain. (**A**) Cartoon representation of the dARC1 CA dimer, the N-terminal extended β-strand and α-helices are numbered sequentially from the N- to the C-terminus. Monomer-A is coloured cyan and Monomer-B in wheat. The right-hand panel is a view at 90° relative to the left-hand panel. (**B**) Close-up cartoon representations of dARC1 CA-NtD (left) and dARC CA-CtD (right) showing the helical topology of each domain. (**C**) 3D Cα structural alignment of dARC1 CA-NtD (blue cartoon) with dARC1 CA-CtD (red cartoon) with secondary structure elements labelled.

### The dARC1 CA dimer interface

The dARC1 CA-CtD monomer consists of a five-helix core comprising α5 (residues A125-Q134), α6 (residues I143-Q156), α7 residues (E164-L171), α8 (residues I177-H182) and α9 (residues F191-N204). The dimer interface is located between CA-CtDs where the outer surfaces of α5 and α7 pack against α5’ and α7’ of the opposing monomer (**Fig. 2A**). The homodimer interface encompasses 1005 Å^2^ of buried surface and is defined by numerous intermolecular interactions. The interface is largely hydrophobic with contributions from sidechain packing of the Y126, Y129, M130, F133, L170, F172 and L174 hydrophobic and aromatic residues that are exposed on α5 and α7 and form a continuous apolar network with a Y129 and F133 at its centre (**Fig. 2A**). This is apparent in the analysis of the dARC1 CA surface hydrophobicity profile reveals a distinct apolar patch that locates to the centre of the CA-CtD homodimer interface (**Fig. S2A**). In addition, at the periphery of the interface, there is also a salt bridge between R161 on the α6-α7 connecting loop with D169 at the C-terminus of α7 providing further stabilisation (**Fig. 2A**). The number and hydrophobic nature of interactions within the homodimer interface suggest the dimer constitutes a relatively stable or obligate structure.

**Fig. 2.**
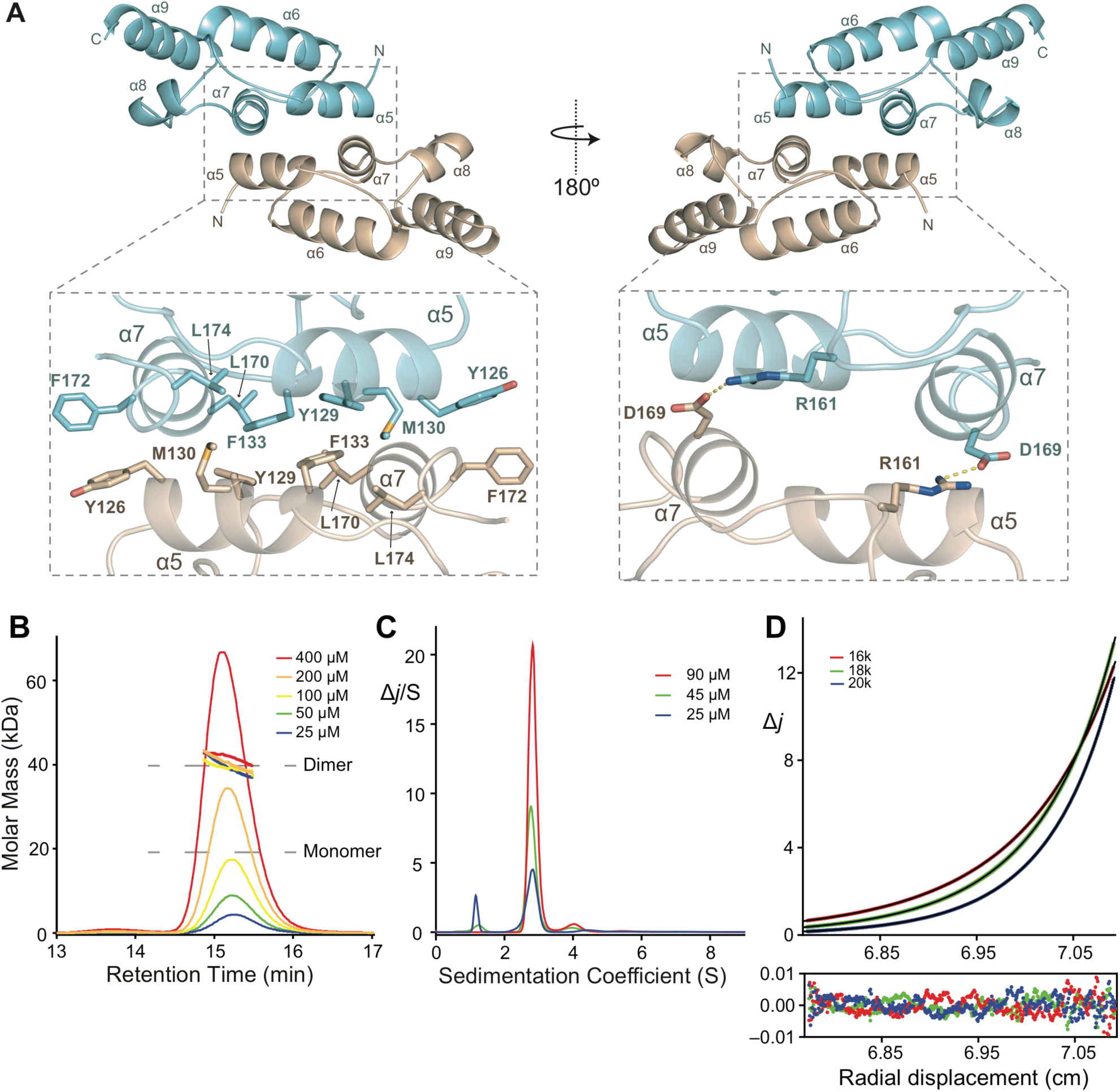
The dARC1 CA dimer interface and solution conformation. (**A**) A cartoon representation of dARC1 CA-CtD dimer, α-helices are numbered sequentially from the N- to the C-terminus. Monomer-A is coloured cyan and Monomer-B in wheat. The right-hand panel is a view at 180° relative to the left-hand panel. **Inset,** close up of molecular details of interactions at the dARC1 dimer interface. Residues that make interactions are shown in stick representation coloured by atom type. Salt-bridge interactions between R161 and D169 are shown as dashed lines. (**B**) SEC-MALLS analysis of dARC1 CA. The sample loading concentrations were 400 µM (8 mg/mL) (red), 200 µM (4 mg/mL) (orange), 100 µM (2 mg/mL) (yellow), 50 µM (1 mg/mL) (green) and 25 µM (0.5 mg/mL) (blue). The differential refractive index (dRI) is plotted against column retention time and the molar mass, determined at 1-second intervals throughout the elution of each peak, is plotted as points. The dARC1 CA monomer and dimer molecular mass is indicated with the grey dashed lines. (**C**) C(S) distributions derived from sedimentation velocity data recorded from dARC1 CA at 25 µM (blue), 50 µM (green) and 100 µM (red). The curves represent the distribution of the sedimentation coefficients that best fit the sedimentation data (ƒ/ƒ_0_ = 1.41). (**D**) Multi-speed sedimentation equilibrium profile determined from interference data collected on dARC1 CA at 70 µM. Data was recorded at the three speeds indicated. The solid lines represent the global best fit to the data using a single species model (Mw = 38.9±1 kD). The lower panel shows the residuals to the fit.

### Self-association of dARC1 CA

Given the unexpected nature of the dimer observed in the crystal structure, the solution molecular mass, conformation and self-association properties of dARC1 CA were examined using a variety of solution hydrodynamic methods. Initial assessment by Size Exclusion Chromatography coupled Multi-Angle Laser Light Scattering (SEC-MALLS) was performed with protein concentrations ranging from 25-400 µM that yielded an invariant solution molecular weight of 40.0 kDa for dARC1 CA (**Fig. 2B**). By comparison, the dARC1 CA sequence-derived molecular weight is 19.6 kDa. Given this value, together with the lack of a concentration dependency of the molecular weight, it is apparent that dARC1 CA also forms strong dimers in solution. To confirm and better analyse dARC1 CA oligomerisation, the hydrodynamic properties were analysed using sedimentation velocity (SV-AUC) and sedimentation equilibrium (SE-AUC) analytical ultracentrifugation. A summary of the experimental parameters, molecular weights derived from the data, and statistics relating to the quality of fits are shown in **Table S2**. Analysis of the sedimentation velocity data for dARC1 CA using both discrete component and the C(S) continuous sedimentation coefficient distribution function **Fig. 2C** revealed a predominant single species with S_20,w_ of 2.92±0.03 S and no significant concentration dependency of the sedimentation coefficient over the range measured (25-90 µM). These data show that dARC1 CA comprises a single stable 2.92 S species with a molecular weight derived from either the C(S) function or discrete component analysis (S_20,w_/D_20,w_) of 38 kD, **Table S2**, consistent with a dARC1 CA dimer. The frictional ratio (f/f_o_) obtained from the analysis of the sedimentation coefficients is 1.41 (**Table S2**), suggesting the solution dimer has an elongated conformation and is consistent with the U-shaped conformation observed in the crystal structures. Moreover, analysis of the crystal structure using HYDROPRO (*21*) gives calculated S_20,w_ and D_20,w_ values, in close agreement with that observed in solution (**Table S2**), supporting the idea that the dimer observed in the crystal structures is wholly representative of the solution conformation. To further ascertain the affinity of dARC1 CA self-association, multispeed SE-AUC studies at varying protein concentration were carried out and typical equilibrium distributions for dARC1 CA are presented in **Fig. 2D**. Analysis of individual gradient profiles showed no concentration dependency of the molecular weight and so all the data were fitted globally with a single ideal molecular species model, producing a weight averaged molecular weight of 38.9 kDa, **Table S2**. The lack of any concentration dependency precludes any analysis of homodimer affinity but confirms that dARC1 CA forms a stable dimeric structure that has the expected properties of the dimer we observe in the crystal structure.

Attempts to mildly disrupt the central apolar network by introduction of an F133A mutation had no effect on dimerization, when assessed by SEC-MALLS (**Fig. S2B**). More aggressive mutations F133A+Y129A and F133A+R161A resulted in complete loss of protein solubility and an inability to purify the constructs, further suggesting that in dARC1 CA homodimerization is requirement for protein folding/structural integrity and likely forms a key building block of dARC1 particle assembly. Analysis of the electrostatic surface potential of the dimeric structure reveals a differential distribution of charge where the surface of the glacial trough has a net negative charge that spreads across both domains of each dARC1 and the underside were the C-terminus projects has a more positively charged character (**Fig. S2C**), suggesting that upon assembly dARC1 particles would have a negatively charged exterior and more positively charged interior were nucleic acid is contained.

### Comparison with mammalian ARC CA structure

Given that mam-ARC and dARC1 share functional similarities, we assessed the relationship between rARC and dARC1 by comparing the dARC1 structure with the individual domains from rARC. Overall, the alignments are excellent, reflecting the evolutionary relationship, but there are significant differences between dARC1 and rARC in both their NTDs and CTDs.

There are two crystal structures of the rARC NTD in complex with peptide ligands (PDB 4X3H and 4X3I) (Zhang et al., 2015), and a recent solution NMR structure (6GSE, Nielsen et al., 2019) of the entire rARC CA domain that resolves the NTD in an apo form. Superficially, the dARC1 CA-NTD aligns well with all available structures of the rARC CA-NTD, with DALI Z-scores and RMSDs of 8-10 and 1.5-1.9 Å respectively (**Fig. 3A**)

**Fig. 3.**
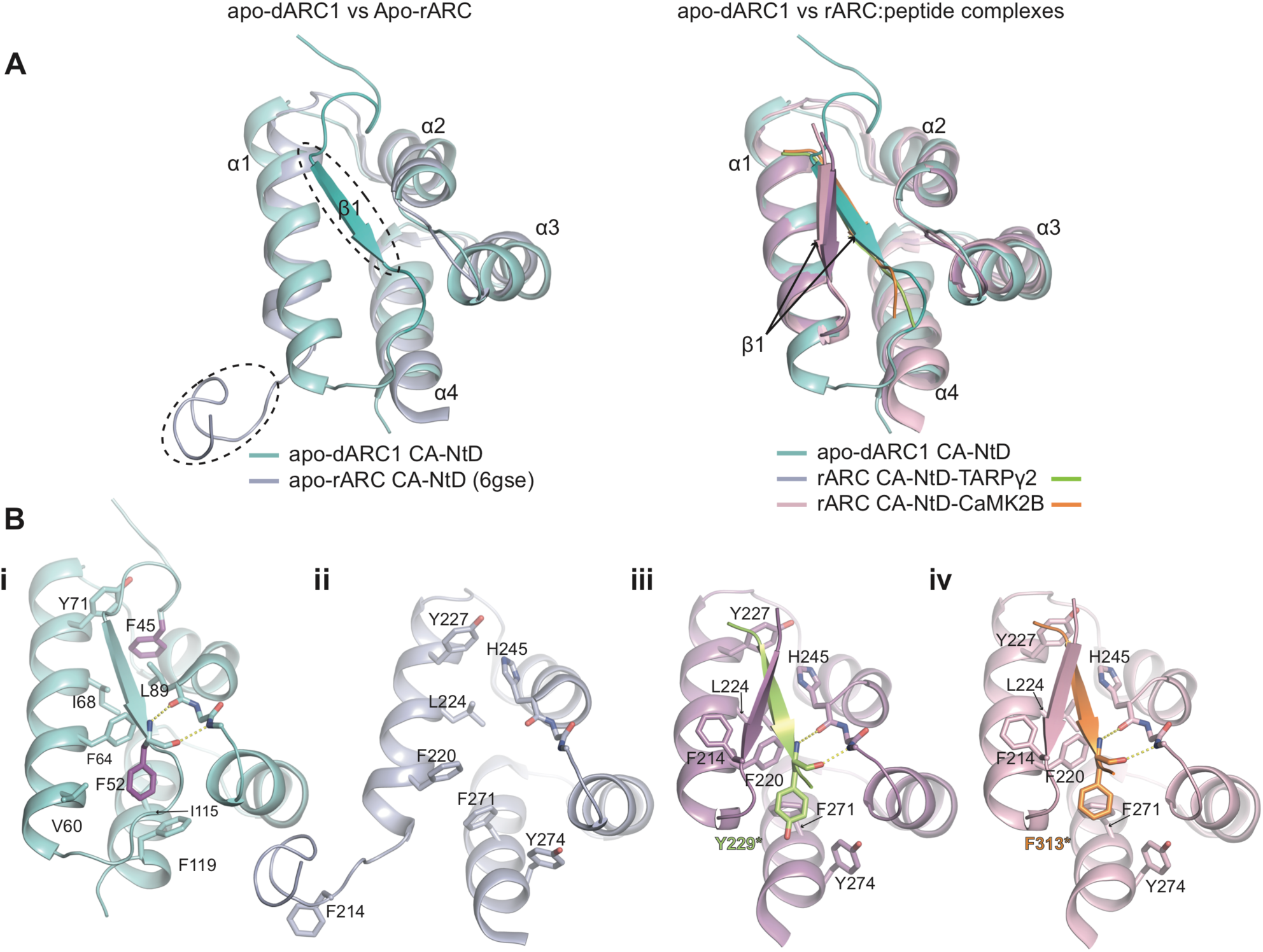
Comparison of dARC1 and rARC CA-NTD structures. (**A**) Left, 3D structural alignment of dARC1 CA-NtD (teal cartoon) and the apo-rARC CA-NtD (PDB: 6GSE, lilac cartoon). Secondary structure elements are labelled. Circled are the ordered, N-terminal β-strand of dARC1, and the disordered N-terminal strand of apo-rARC. Right, 3D structural alignment of dARC1 CA-NtD and the peptide-complex structures of rARC CA-NtDs (PDB: 4X3H and 4X3I). The protein backbones are shown in cartoon representation coloured according to the legend. Secondary structure elements are labelled. The arrow indicates the different positioning of the extended N-terminal β-strand between the dARC1 and rARC structures. (**B, i to iv**) Individual views of the structures presented in **A**. **i**) apo-dARC1, **ii**) apo-rARc, **iii**) rARC-TARPγ2, **iv**) rARC-CaMK2B. Residues that constitute the hydrophobic NtD cleft are shown in stick format, coloured by atom type. In each structure, the side chain of the aromatic residues buried in the interface (F45 and F52, dARC1 CA-NtD; Y229*, rARC CA-NtD-TARPγ2; F313*, rARC CA-NtD-CaMK2B) are coloured purple, yellow and orange respectively. The conserved mainchain hydrogen bonding interactions between the backbone amide and carbonyl of F52 with the carbonyl of L89 and the amide of Y91 (dARC1), of Y229 with the carbonyl of H245 and the amide of N247 (rARC CA-NtD-TARPγ2) and of F313 (rARC CA NtD-CaMK2B) with the carbonyl of H245 and the amide of N247 are shown as dashed lines.

Examination of the dARC1 CA-NtD reveals an N-terminal extended strand (NT-strand), residues G43 to R56, with a short β-configuration that packs against the core of the NtD. The NT-strand makes many interactions with the apolar and aromatic side chains that extend from α1, α2 and α4, burying 803 Å^2^ of surface in the interface (**Fig. 3A and Bi, Fig. S3A**) and the same configuration is observed in all four instances of the NtDs that we see in our two crystal structures (**Fig. S3B**). The NT-strand residues are highly conserved in *dARC* genes across Drosophilidae but not with the Mam-ARCs (**Fig. S3C**). In particular, two highly conserved aromatic residues, F45 and F52, are entirely buried, surrounded by the conserved side chains of F64, L89, I115, and F119 and act to anchor the NT-strand into the hydrophobic α1-α4 cleft of the CA-NtD. In addition, there is a mainchain interaction between the backbone amide and carbonyl of F52 with the carbonyl of L89 and the amide of Y91 that further stabilises the conformation of the NT-strand (**Fig. 3Bi and Fig. S3A**).

In apo-rARC CA-NTD (6GSE) the helical core aligns very well with the corresponding region of dARC1 (RMSD = 1.45 Å). However, here the rARC NT-strand, residues D210-E216, has a disordered conformation (**Fig. 3, A and Bii**) and the α1-α4 hydrophobic cleft, which in dARC1 contains the native NT-strand, is unoccupied in rARC suggesting there is a functional divergence for the NT-strand between the dARC1 and mam-ARC families. This notion is supported by inspection of the rARC CA NtD-TARPγ2 and CA NtD-CaMK2B complexes (4X3H and 4X3I) where the α1-α4 cleft of rARC is now occupied by the bound TARPγ2 or CaMK2B derived peptides (**Fig 3B iii and iv**) and the bound peptides adopt the same extended β-configuration as the native NT-strand in the dARC1 structure (**Fig. S3D**) and bury a comparable amount of surface, 772 Å and 641 Å, respectively. Moreover, both bound peptides contain an aromatic residue equivalent to dARC1 F52, Y229 in TARPγ2 and F313 in CaMK2B, that packs into the core of rARC CA-NtD and makes an identical main-chain interaction with the backbone carbonyl of H245 and the amide of N247 as that observed between the backbone amide and carbonyl of F52 with the carbonyl of L89 and the amide of Y91 in dARC1 (**Fig. S3D**). In these peptide-complex structures, the rARC NT-strand, D210-E216, that is disordered in the apo-structure now adopts a parallel β-configuration to pack against the bound peptides (**Fig. 3, Biii and Biv**) and it is possible that the propensity to form this stabilising beta configuration has been selected for. This notion is supported by inspection of the dARC and mam-Arc multiple sequence alignment (**Fig. S3C**) that reveals a conserved “TQIF” motif in Amniota that retains β-branched residues, favoured in beta structure, at the T and I position. This motif is not present in amphibians or in *L. ch* Gypsy2 the closest known relative to the transposon from which tetrapod ARC was exapted, suggesting that this feature, and possibly peptide binding ability, arose within Amniota.

The structures of dARC1 CA-CtD and rARC CA-CtD (PDB: 4X3X) also superimpose well (RMSD = 2.7 Å). However, the CtD of the apo-rARC CA NMR structure more closely matched the structure of dARC1 CA-CtD, (RMSD = 2.2 Å), with all 5 helices overlaying (**Fig. 4A**). However, in contrast to our solution studies of dARC1 (**Fig. 2, A to C and Fig S2A**) the rARC CA domain was monomeric in solution, even at the high concentrations under which NMR was performed (*20*).

**Fig. 4.**
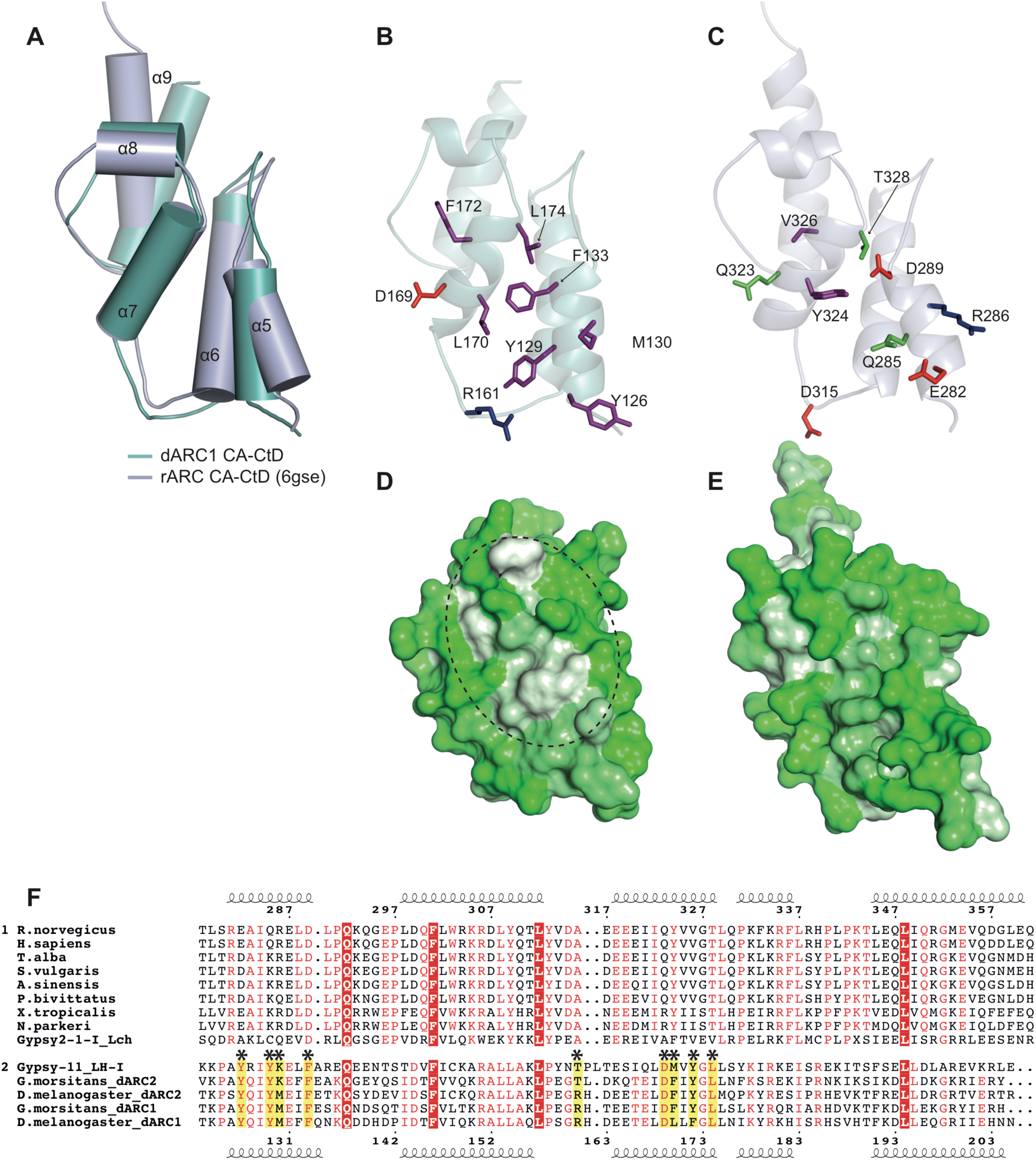
Comparison of the dARC1 and rARC CA-CTD structures. (**A**) 3D structural alignment of dARC1 CA-CtD and the rARC CA-CtD from apo rARC (PDB: 6GSE). The structures are shown in cartoon with equivalent helices labelled and shown as cylinders. dARC1 is coloured cyan and rARC is coloured light blue. (**B and C**) Details of the CA-CtD homodimer interfaces. Cartoon representations of the protein backbone of dARC1 CA-CtD (**B**) and rARC CA-CtD (**C**) are shown, coloured as in **A**. The view is of one monomer looking into the dimer interface, Residues that make interactions in dARC1 CA and their equivalents in rARC are shown in stick representation colour-coded by residue type (purple, hydrophobic/aromatic; green, polar; red, acidic; blue, basic). (**D and E**) Hydrophobic surface representations of **B** and **C** respectively. Circled in **D** is a distinct hydrophobic patch on the surface of the dARC1 CA-CtD which is absent in rARC. (**F**) Multiple sequence alignment of ARC, dARC1 and dARC2 CA-CtDs, and parent retrotransposon sequences. Group 1 contains tetrapod ARC (tARC) sequences and the closely related *Latimeria chalumnae* (*L. ch*). Above, secondary structure of rARC, numbers according to the rARC (*R. norvegicus*) sequence. Group 2 contains dARC1, dARC2 and closely related *Linepithema humile* (*L. h*) Gypsy11 retrotransposon. Below, secondary structure of dARC1, numbers according to the dARC1 (*D. melanogaster*) sequence). Red box, white text, invariant residues shared between groups. Red text, residues conserved within a group. Asterisks mark the residues at the dARC1 CtD dimer interface and their equivalents in tARCs, as shown in **B** and **C**.

In dARC1, a large proportion of the CTD dimer interface results from the packing of hydrophobic side chains projecting from helices 5 and 7 (**Fig. 2A**). However, upon comparison of external α5/α7 surfaces of dARC1 and rARC (**Fig. 4, B and C**) it is apparent that the exposed Y126, Y129, M130, F133, L170, F172 and L174 side chains that are responsible for the hydrophobic character of the dARC1 dimer interface are not conserved in rARC and are replaced by E282, Q285, R286, D289, Y324, V326 and T328 in rARC. Indeed, the hydrophobic patch present on the surface of dARC1 is not evident in the same surface on rARC (**Fig. 4, D and E**). In addition, R161 and D169, which make a salt bridge interaction in the dARC1 interface, are also not conserved, being replaced by D315 and Q323 in rARC (**Fig. 4, B and C**). These sequence differences are also apparent throughout the entire dARC and mam-ARC families. Here, there is strong sequence conservation of residues that constitute the core fold of the CA-CtD across both dARC and mam-ARCs but the hydrophobic CA-CtD dimer interface residues are only present in the dARC lineage (**Fig. 5F**). Taken together, these data reveal that, while tertiary structure topology of dARC1 and rARC CA-CtDs is conserved, there are significant differences in the character of the surface that is presented around α5-α7; in dARC1, the hydrophobic nature of this surface drives the formation of a strong CTD dimer, whereas in rARC, the more polar nature of this surface may explain why the protein is monomeric in solution. Given these differences, although there is strong evidence for assembly of both dARC1 and mam-ARC into capsid-like particles (*12, 15*), it seems likely that if dARC1 and mam-ARC use the α5/α7 interface in a particle assembly pathway, the interface may be substantially weaker for mam-ARC.

**Fig. 5.**
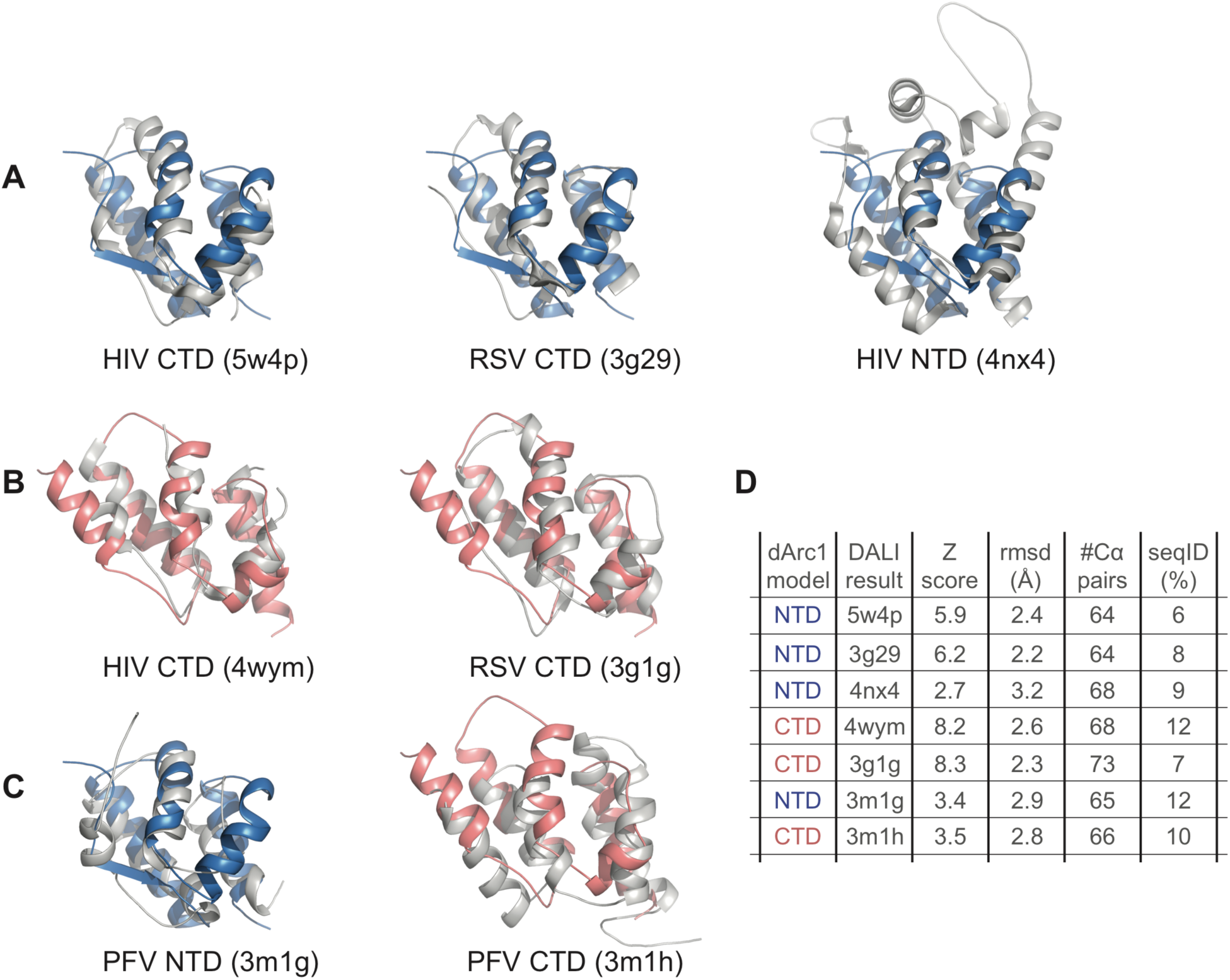
Structural similarity with ortho and spumaretroviral CA. (**A**) Pairwise DALI 3D Cα structural alignment of dARC1 CA-NtD with HIV CA-CtD (Left), RSV CA-CtD (middle) and HIV-NtD (right). In each panel, the cartoon of the dARC1 CA-NtD backbone is shown in blue, the backbone of the aligned structures is shown in grey. (**B**) Pairwise 3D Cα structural alignment of dARC1 CA-CtD with HIV CA-CtD (Left) and RSV CA-CTD (right). In each panel the cartoon of the dARC1 CA-CtD backbone is shown in red, the backbone of the aligned structures is shown in grey. (**C**) Pairwise 3D Cα structural alignment of dARC1 CA-NtD with Prototypic Foamy virus (PFV) CA-NtD (Left) and dARC1 CA-CtD with PFV CA-CTD (right). (**D**) DALI Z scores, RMSD, number of aligned residues and sequence identities for 3D Cα alignments.

### Retroviral CA domain structures are related to both dARC1 CA domains

The topology of the alpha-helical two-domain fold of dARC1 is highly reminiscent of retroviral CA structures. Interrogation of the PDB database with dARC1 CA using the DALI alignment/search engine (*22*) produced an overwhelming number of matches to Gag proteins, (87%, Z-score ≥ 5.0) and identified rARC, together with many orthoretroviral and spumaretroviral CA-NtD and CA-CtD structures. Alignments with CA-NtDs and CA-CtDs from HIV CA, Rous Sarcoma virus (RSV) CA and prototypic foamy virus (PFV) CA are presented in **Fig. 5**. The best structural alignments to dARC1-NtD were with retroviral Gag CA-CtD structures rather than Gag CA-NtD structures (**Fig. 5, A and D**) indicating that the dARC1 CA-NtD is more closely related to the orthoretroviral CA-CtD than it is to the orthoretroviral CA-NtD. Alignments with dARC1-CtD also had the best structural alignment with orthoretroviral Gag CA-CtD structures (**Fig. 5, B and D**), perhaps not surprising given the observation of close resemblance of the dARC1 CA-NtD to the dARC1 CA-CtD (**Fig. 1Biii**). Alignments with PFV CA-NtD and CA-CtD were also found (**Fig. 5C and D**) and although not as significant as with the orthoretroviral CA, these data support previous observations of a relationship of spumaretroviral Gag with mam-ARC (*17*).

These data provide evidence for a structural conservation between orthoretroviral CA and ARC proteins and the weaker alignments observed with orthoretroviral CA-NtDs suggest that orthoretroviral CA-NtDs have undergone much more structural divergence than has occurred in the Ty3 family or ARC proteins. Moreover, these data further support the previously proposed idea that a duplication of a CA-CtD progenitor first gave rise to double domain ancestors and that subsequent divergence of domains resulted in spumaretroviral, orthoretroviral, and Metaviridae-derived proteins such as ARC, that are found presently (*17, 23*).

### The dARC1 CtD dimer is an ancient assembly interface conserved in orthoretroviridae

Given the existence of the dARC1 CA dimer and the distant relationship with orthoretroviral CA, we next looked to see if the dimer interface was conserved between dARC1 and the CtD dimers of HIV-1 CA and RSV CA that are known to be essential for capsid assembly in orthoretroviruses. Cartoon representations of the dARC1, HIV-1 and RSV CA-CtD dimers are shown in **Fig. 6, A to C**. In each, the domain arrangement that presents the dimer interface is the same and this also seen in the CA-CtD dimer of native Ty3 particles visualised by cryo-electron microscopy (*24*) but with some repositioning of the CA-NtDs (**Fig. S4**). The structures have been aligned to find the best Cα alignment over the entire dimer (HIV, RMSD = 2.8 Å over 117 Cα; RSV, RMSD = 3.1 Å over 101 Cα) (**Fig. 6, D and E**) and it is apparent that each interface is made up from interactions between residues on CtD helices α5 and α7 of dARC1, that correspond to α7’ and α8 in the orthoretroviral CA-CtD structures. Notably in the orthoretroviruses, α7’ is reduced to a single turn and the monomers are rotated with respect to each other. Therefore, in dARC1 residues on α5 and α7 contribute equally to the interface whilst in the orthoretroviruses α8 contributes more to the interface than α7’. This combination of the larger contribution of α5 in dARC1, together with the rotation and displacement of CA-CtDs seen in the orthoretroviruses, has the effect of reducing the surface area that is buried at the interface from 768 Å^2^ in dARC1 to 452 Å^2^ in HIV-1. Notably, the homodimer affinity for orthoretroviral CA-CtD dimers is much weaker than the dARC1 dimer. Equilibrium dissociation constants ranging between 10-20 µM have been reported for HIV-1 (*25, 26*) and CA-CTD dimerisation is undetectable for other genera (*27–29*). Nevertheless, given the domain organisation and the similarity in character of the orthoretroviral and dARC1 CA-CtD dimers, we suggest this interface is a key building block of capsid assembly, retained in dARC1 and conserved from Ty3/Gypsy transposable elements through to orthoretroviridae.

**Fig. 6.**
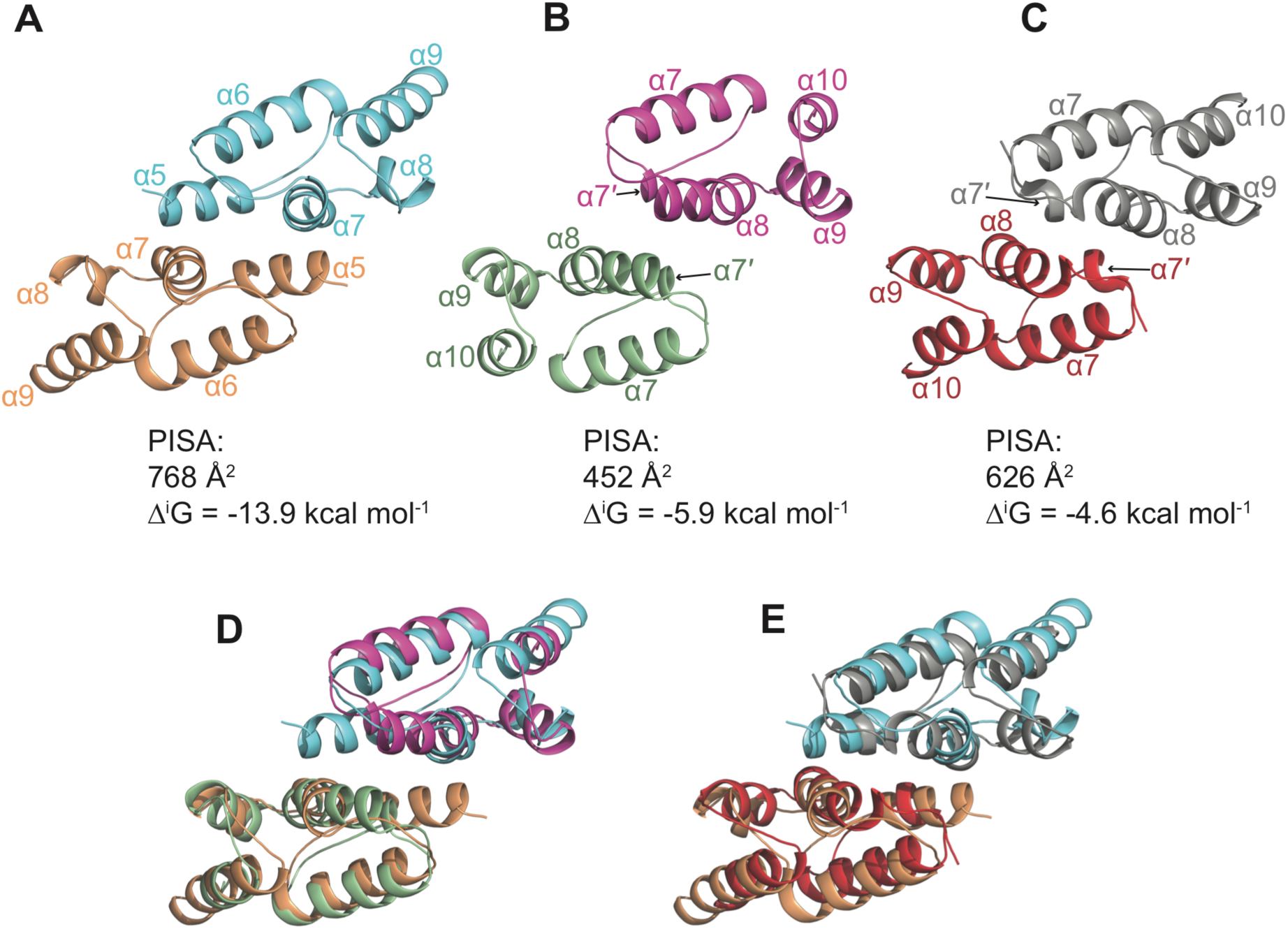
Comparison with retroviral CA-CtD dimers. **(A to C)** Cartoon representations of CA-CtD dimers. (**A**) dARC1, cyan and wheat, (**B**) HIV-1, magenta and pale green, (**C**) RSV, grey and red. The orthoretroviral structures are aligned with respect to the dARC1 dimer. CtD helices α5 to α9 are labelled in the dARC1 structure and the equivalent α7’ to α10 in the orthoretroviral structures. The buried surface area (Å^2^) and free energy of interaction (Δ^i^G) of each interface, calculated in PDBePISA is displayed below each structure. (**D and E**) Structural alignment of dARC CA with HIV-1 CA and RSV CA dimers respectively. Protein backbones are coloured as in **A, to C**.

## DISCUSSION

### dARC1 Capsid structures

Our crystal structures demonstrate that the central region of dARC1 contains two largely α-helical domains that, despite the lack of sequence conservation, have the same predominantly α-helical folds observed in the structures of CA domains from the ortho- and spuma-retroviruses. A more detailed inspection of dARC1 CA-NtD and CA-CtD reveal they comprise 4- and 5-helix bundles, respectively, with a topology that aligns well with the arrangement of secondary structure elements observed in orthoretroviral CA NtDs and CtDs (**Fig. 5**). However, it is apparent that both the ARC CA-NtD and CA-CtD are much more closely related to the orthoretroviral CA-CtDs than they are to orthoretroviral CA-NtDs (**Fig. 5**), consistent with our previous notion that an ancient domain duplication was a key event during retrotransposon evolution (*17*). Notably, orthoretroviral CA-NtDs contain an extra N-terminal β-hairpin and an additional two helices compared to the ARCs and the CA domains of Ty3/Gypsy transposons (*24*) (**Fig. S4**). This suggests that unique aspects of the retroviral life cycle might be driving specific changes in structure of the retroviral CA-NtD. One such pressure might be associated with the process of maturation that follows retrovirus budding from the cell. Maturation involves proteolytic cleavage of immature viral cores followed by CA reassembly to yield mature virions and although it is proposed that dARC1 and mam-ARC transport mRNA between cells it is thought likely that particles are packaged into extracellular vesicles for cell-to-cell transfer (*12, 15*). Similarly, maturation events do not occur in Ty3 elements, that also do not bud from the cell, and that have Gag which assembles directly into mature forms (*24*). Absence of maturation also characterizes spumaviruses and it was observed previously that the CA NtD-equivalent region of PFV Gag showed greater similarity to rARC than to orthoretroviral CA (*17*).

### Structural differences between insect and mammalian ARCs

Our 3D superimpositions have demonstrated that there is a large degree of structural conservation between the dARC1 and mam-ARC CA structures. However, despite this strong similarity, two regions of distinct differences between the dARC1 and rARC structure are apparent. The first concerns the ARC CA-NtD and the interaction with potential binding partners, the second, the putative dimerization domain of the CtD.

Functionally important interactions between mam-ARC and a variety of neuronal proteins, including the TARPγ2 and CaMK2B proteins, as well as the NMDA receptor, have been defined (*16, 20*). However, no such interactions have been reported for dARC1. In the rARC structures with bound TARPγ2 or CaMK2B peptides the disordered N-terminal region of rARC seen in the apo structure, now forms a short parallel β-sheet with the bound peptide stabilising the peptide binding within a hydrophobic cleft on rARC. It is apparent that the conformation of these rARC-bound peptides strongly resembles the conformation of the NT-strand of dARC1 NtD (**Fig. 3**). Therefore, given the sequence differences in the NT-strand region between the dARC and mam-ARCs (**Fig. S3C**) one notion is that mam-ARC has evolved an N-terminal strand that no longer binds into the CA-NtD hydrophobic cleft but has gained the ability to promote the binding of synaptic protein ligands, perhaps acting as a sensor of synaptic stimuli. This sensing property might then contribute control to a functional role for ARC based on assembly and mRNA trafficking.

There are also significant differences between the dARC1 and rARC CA-CtD, illustrated in **Fig. 4**. Overall, our crystal structure of dARC1 and the NMR structure of full length rARC (Nielsen et al., 2019) are very similar, with good overlay in all five helices. However, inspection of the dARC1 surface reveals a substantial hydrophobic patch absent in rARC (**Fig. 4D and E**). This hydrophobic patch is shared with the orthoretroviruses (*25, 30*) and seems to be associated with the formation of stable dARC1 dimers, whereas rARC is monomeric. Whether this translates to differences in the stability of assembled particles *in vivo* remains to be determined, however it is possible that differences in the physiological roles of dARC1 and mam-ARC may mean that mam-ARC has evolved to require a weaker interface that facilitates disassembly. Alternatively, it is possible that mam-ARC may require a conformational change to facilitate dimerization or uses a completely different assembly mechanism that employs other surfaces of the molecule.

### ARC particle assembly

The observation that residues at the dARC1 CA-CtD interface are not conserved between the insect and mammalian ARC lineages suggests the possibility that, although mam-ARC particles have been observed *in vitro* and in cells, their mode of assembly may not utilise an obligate CA-CtD dimer as a building block. This type of observation has been made with orthoretroviruses that assemble through a combination of NtD-NtD, NtD-CtD and CtD-CtD interactions to form the viral capsid, where the relative contribution that different types of CA interaction make to the overall formation of the viral core varies depending on the retroviral genera. For instance, in lentiviruses, it is apparent that capsid assembly requires a strong intrinsic CtD-CtD dimeric interaction (*25, 30*). However, more generally, capsid shell formation requires three types of interaction, intra-hexamer NtD-NtD self-association (*30–33*), intrahexamer NtD-CtD interactions between adjacent capsid monomers (*30, 34, 35*) and inter-hexamer CtD-CtD interactions (*25, 30*). Therefore, it is entirely possible that in dARC1 and mam-ARC particles the relative contribution of each type of interfaces may also differ.

### ARC exaptation

Mam-ARC and dARC1 appear to have different biological properties. However, it remains to be determined whether these differences result from the capture of two different Ty3/Gypsy elements or whether they reflect evolutionary adaptations. Perhaps the best studied example of the appropriation of retroelement encoded genes by mammalian hosts is the case of syncytin, a fusagenic protein essential for proper placenta formation (*36*). It is evident that syncytin capture appears to have occurred on multiple independent occasions, involving envelope proteins from clearly different retroviruses (*37, 38*), resulting in placentae with subtly different morphologies (*39*). Determining whether this is also the case with the ARC genes, as well as their close relatives in the mammalian genome (*14*) will require further characterization of existing retrotransposon elements utilizing structural methods not reliant on the comparative similarities in related nucleic acid sequences that have disappeared with the passage of time.

## MATERIALS AND METHODS

### Protein expression and purification

dARC1 residues S39-N205 were determined to represent the CA domain according to MSA and secondary structural analysis performed in ClustalX (*40*) and Pspired (*41*). An *E. coli* codon-optimised cDNA for *Drosophila melanogaster* dARC1 (Uniprot Q7K1U0) was synthesised (Geneart), the relevant sequence PCR-amplified, and subcloned into a pET22b plasmid (Novagen). The resulting construct comprises residues 39-205 of dARC1, with an N-terminal Met, and a C-terminal PLEHHHHHH motif. Proteins were expressed in *E. coli* strain BL21 (DE3) grown in LB broth, by induction of log-phase cultures with 1 mM IPTG, incubated overnight at 20°C. Cells were pelleted and resuspended in 50 mM Tris-HCl, 150 mM NaCl, 10 mM Imidazole, 5mM MgCl_2_, 1mM DTT, pH 8.0, supplemented with 1 mg/mL lysozyme (Sigma-Aldrich), 10 µg/mL DNase I (Sigma-Aldrich) and 1 Protease Inhibitor cocktail tablet (EDTA free, Pierce) per 40 mL of buffer. Cells were lysed using an EmulsiFlex-C5 homogeniser (Avestin) and dARC1 CA captured from clarified lysate using immobilised metal ion affinity on a 5 mL Ni^2+^-NTA superflow column (Qiagen). Bound dARC1 CA was eluted in non-reducing buffer (50 mM Tris-HCl, 150 mM NaCl, 300 mM Imidazole) and Carboxypeptidase A (CPA, Sigma C9268) was added at a ratio of ∼ 100 mg dARC1 per mg CPA. The resulting mixture was incubated overnight at 4 °C to allow digestion of the C-terminal His-tag. The CPA was inactivated via the addition of TCEP to 2mM. dARC1 CA was further purified by size exclusion chromatography using a Superdex 75 (26/60) (GE healthcare) column, equilibrated in 20 mM Tris-HCl, 150 mM NaCl, 1mM TCEP pH 8.0. Purified protein eluted in a single peak. Seleno-methionine derivative protein was produced using an identical procedure, although *E. coli* B834 (DE3) cells, grown in seleno-methionine Medium (Molecular Dimensions, Newmarket, UK), were used to express the protein. Electrospray-ionisation mass spectrometry was used to confirm the identity of dARC1, and where applicable, seleno-methionine incorporation. It also confirmed that the N-terminal Met had been processed, and that the His-tag had been completely digested, leaving the motif “PLE” at the C-terminus. Protein was concentrated by centrifugal ultrafiltration (Vivaspin (MWCO 10 kDa), then snap frozen and stored at - 80°C. Concentrations were determined by UV absorbance spectroscopy using an extinction coefficient at 280 nm.

### Protein crystallisation and structure determination

dARC1 CA was crystallised using sitting drop vapour diffusion at 18 °C, using Swissci MRC 2-drop trays (Molecular Dimensions) with drops set using a Dragonfly robot with humidity chamber (TTP Labtech). Native protein was initially concentrated to 20 mg/mL. Typically drops were 200-300 nL, made by mixing protein:mother liquor in a 3:1 or 1:1 ratio, with a 75 µL reservoir. Initial crystal hits were obtained using the Structure Screen 1&2 (Molecular Dimensions) in a condition containing 4.3 M NaCl, 0.1 M HEPES pH 7.5. Two crystal forms could be observed in such conditions. Thin rods which turned out to be oP or as hexagonal disks or trapezoidal prisms which were hP. Datasets were collected for these native crystals, but they could not be solved by molecular replacement methods. SeMet dARC1 CA was crystallised in conditions that optimised protein concentration, NaCl concentration, and pH. The best crystals grew in 300-400 nL drops set with protein at 12.5-16 mg/mL, with mother liquor NaCl ranging between 2.8-3.3 M. Rods were ∼400×30×30 µm and hexagons/trapezoids were ∼130 µm across and up to 30 µm thick. Crystals were harvested using MiteGen lithographic loops. The best cryoprotection was achieved using sodium malonate, mixed into mother liquor to a concentration of 1.6 M. This was added directly to the drop, or crystals were bathed in this solution before flash freezing in liquid nitrogen.

### Data collection and structure determination

Data were collected at the tuneable SLS beamline, PXIII. For the orthorhombic crystal form, a peak dataset was collected to 2.06 Å (see Table S1). The data were processed by the SLS GoPy pipeline in P2_1_2_1_2_1_, using XDS (*42*) and showed significant anomalous signal to 2.82 Å. The resultant dataset was solved using SAD methods with Phenix (*43*) and despite a relatively low FOM the experimental map was readily interpretable and it was possible to almost completely autobuild an initial structure with BUCCANEER (*44*). A higher resolution (1.55 Å) dataset was collected at a non-anomalous, low energy remote wavelength (Table S1). This dataset was processed using the Xia2 (*45*) pipeline, using DIALS (*46*) for indexing and integration, and AIMLESS (*47*) or scaling and merging. This dataset was initially used for refinement to 1.7 Å and manual model building in COOT (*48*). It was evident that the data were anisotropic, and that they might benefit from anisotropic correction. Diffraction images were reprocessed using the autoPROC pipeline (*49*), using XDS, POINTLESS (*50*), AIMLESS and STARANISO (http://staraniso.globalphasing.org/cgi-bin/staraniso.cgi). This dataset was used for further refinement of the model and indeed there was an improvement in map quality, and in agreement between model and data. For the hexagonal crystal form, a highly redundant peak dataset was collected to 2.14 Å. This was processed using the Xia2 pipeline, using DIALS for indexing and integration, and AIMLESS for scaling and merging, showing significant anomalous signal to 2.59 Å, in P6_1_22. This dataset was solved using SAD methods in Phenix. Again, the experimental map was readily interpretable and it was possible to almost completely autobuild an initial structure with BUCCANEER. Refinement and model building were carried out in Phenix and COOT respectively. Anomalous signal was very strong in this dataset and so Freidel pairs were treated separately during refinement. Molprobity (*51*) and PDB_REDO (*52*) were used to monitor and assess model geometry. Details of data collection, phasing and structure refinement statistics are presented in **Table S1**.

### SEC-MALLS

Size exclusion chromatography coupled multi-angle laser light scattering (SEC-MALLS) was used to determine the molar mass of dARC CA. Samples ranging from 25 µM to 400 µM were applied in a volume of 100 µL to a Superdex INCREASE 200 10/300 GL column equilibrated in 20 mM Tris-HCl, 150 mM NaCl and 0.5 mM TCEP, 3 mM NaN_3_, pH 8.0, at a flow rate of 1.0 mL/min. The scattered light intensity and the protein concentration of the column eluate were recorded using a DAWN-HELEOS laser photometer and OPTILAB-rEX differential refractometer respectively. The weight-averaged molecular mass of material contained in chromatographic peaks was determined from the combined data from both detectors using the ASTRA software version 6.0.3 (Wyatt Technology Corp., Santa Barbara, CA, USA).

### Analytical Ultracentrifugation

Sedimentation velocity experiments were performed in a Beckman Optima Xl-I analytical ultracentrifuge using conventional aluminium double sector centrepieces and sapphire windows. Solvent density and the protein partial specific volumes were determined as described (*53*). Prior to centrifugation, dARC1 CA samples were prepared by exhaustive dialysis against the buffer blank solution, 20 mM Tris-HCl pH 8, 150 mM NaCl and 0.5 mM TCEP (Tris Buffer). Samples (420 µL) and buffer blanks (426 µL) were loaded into the cells and centrifugation was performed at 50,000 rpm and 293 K in an An50-Ti rotor. Interference data were acquired at time intervals of 180 s at varying sample concentration (25, 50 and 100 µM). Data recorded from moving boundaries was analysed in terms of the size distribution functions C(S) using the program SEDFIT (*54–56*).

Sedimentation equilibrium experiments were performed in a Beckman Optima XL-I analytical ultracentrifuge using aluminium double sector centrepieces in an An-50 Ti rotor. Prior to centrifugation, samples were dialyzed exhaustively against the buffer blank (Tris Buffer). Samples (150 µL) and buffer blanks (160 µL) were loaded into the cells an after centrifugation for 30 hours, interference data was collected at 2 hourly intervals until no further change in the profiles was observed. The rotor speed was then increased and the procedure repeated. Data were collected on samples of different concentrations of dARC1 CA (25, 50 and 70 µM) at three speeds and the program SEDPHAT (*57, 58*) was used to determine weight-averaged molecular masses by nonlinear fitting of individual multi-speed equilibrium profiles to a single-species ideal solution model. Inspection of these data revealed that the molecular mass of dARC1 CA showed no significant concentration dependency and so global fitting incorporating the data from multiple speeds and multiple sample concentrations was applied to extract a final weight-averaged molecular mass.

### Structure analysis and alignments

Molecular interfaces were analysed using the EBI protein structure interface analysis service PDBePISA (https://www.ebi.ac.uk/pdbe/pisa). Electrostatic surface potential and the surface hydrophobicity/hydrophilicity distribution of the dARC1 CA dimer were calculated with APBS (*59*) and using the pymol script (https://pymolwiki.org/index.php/Color_h) respectively. The DALI (*60*) comparison server (http://ekhidna2.biocenter.helsinki.fi/dali) was used to search for and align structural homologues from the PDB.

### Sequence alignment

Amino acid alignments were produced with MAFFT v7.271 (*61*), within tcoffee v11.00.8cbe486 (*62*), weighting alignments using 3-state secondary-structure predictions produced with RaptorX Property v1.02 (*63*). Alignment images were produced with ESPript (*64*).

## Supporting information

Cottee_et_al_6S7Y_PDB_report

Cottee_et_al_6S7X_PDB_report

## General

We thank the Swiss Light Source for beamtime, and the staff of beamline PXIII.

## Author contributions

M.A.C., S.C.L and I. A. T. performed experiments. M.A.C., S.C.L, G.R.Y., J.P.S. and I.A.T. contributed to experimental design, data analysis and manuscript writing.

## Data and materials availability

The coordinates and structure factors for dARC1 CA (S39-N205) have been deposited in the Protein Data Bank under accession numbers 6S7X and 6S7Y.

## Supplementary Materials

**Table S1.**
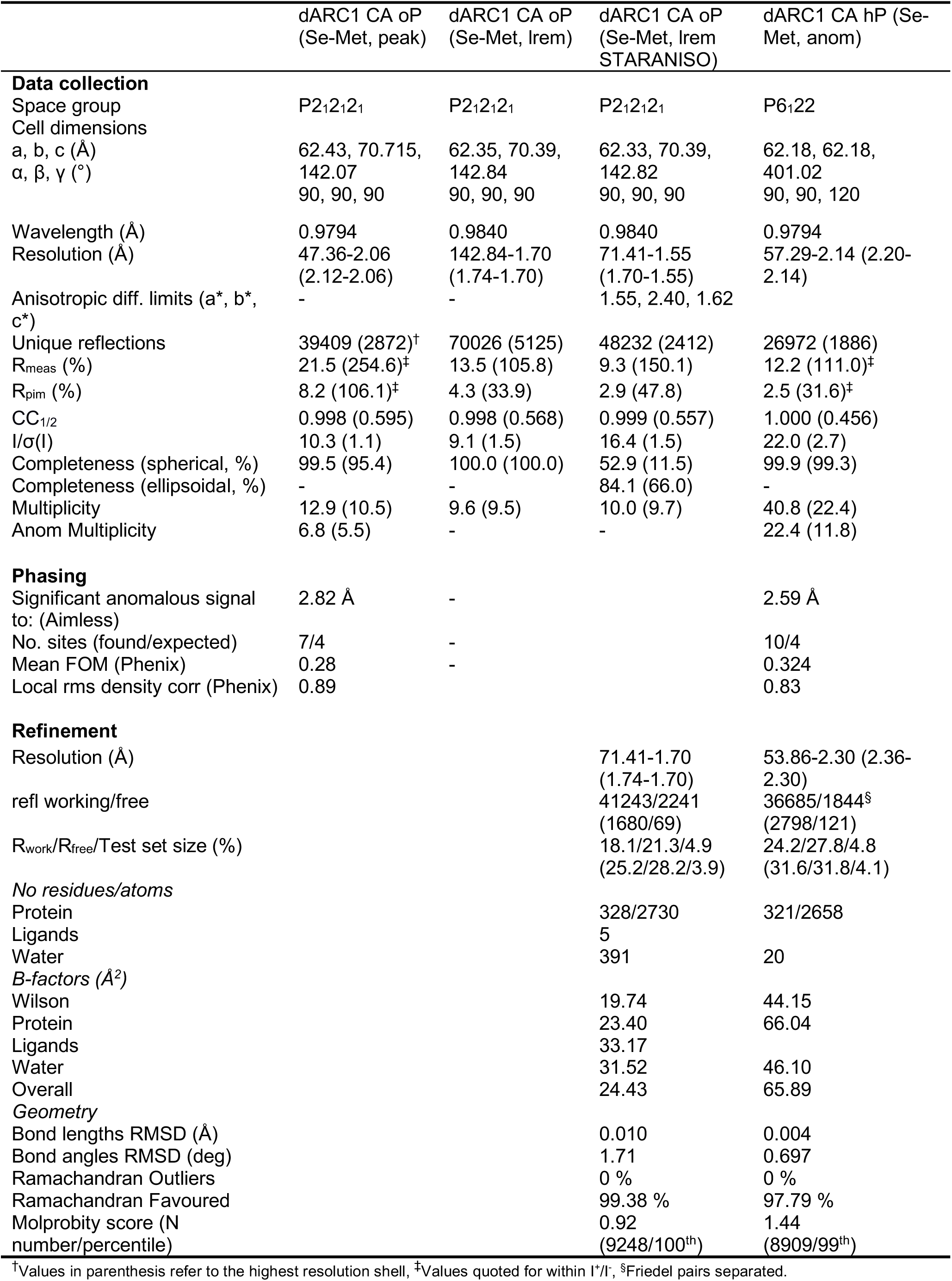
dARC1 CA Statistics of data collection, phasing and refinement.

**Table S2.** Hydrodynamic parameters of dARC1 CA.

| Protein | dARC1 CA |
| --- | --- |
| **Protein Parameter** |  |
| $v$ (mL.g<sup>-1</sup>) | 0.731 |
| $\rho$ (g.mL<sup>-1</sup>) | 1.005 |
| <sup>a</sup>$M_r$ | 19,633 |
| $\epsilon_{280}$ (M<sup>-1</sup> cm<sup>-1</sup>) | 23,600 |
| **Sed velocity** |  |
| $C_{range}$ (μM) | 25-100 |
| <sup>b</sup>$S_{20,w}$ (x10<sup>13</sup>) sec | 2.92±0.03 |
| <sup>c</sup>$S_{20,w}$ (calc) (x10<sup>13</sup>) sec | 2.93 |
| <sup>d</sup>$D_{20,w}$ (x10<sup>7</sup>) cm<sup>2</sup> sec<sup>-1</sup> | 6.95±0.05 |
| <sup>e</sup>$D_{20,w}$ (calc) (x10<sup>7</sup>) cm<sup>2</sup> sec<sup>-1</sup> | 6.77 |
| <sup>f</sup>$M_w$ (S/D) kD | 38.2±1 |
| <sup>g</sup>$f/f_0$ ($S_{20,w}$) | 1.41 |
| <sup>h</sup>$f/f_0$ ($D_{20,w}$) | 1.37 |
| <sup>i</sup>$M_w$ C(S) kD | 38.7±3 |
| <sup>j</sup>$f/f_0$ C(S) | 1.41 |
| <sup>k</sup>RMSD C(S) | 0.005 – 0.008 |
| **Sed eqm** |  |
| $C_{range}$ (μM) | 25-70 |
| <sup>l</sup>$M_w$ kD | 38.9±1 |
| <sup>m</sup>RMSD | 0.002 – 0.004 |
| <sup>n</sup>$\chi^2$ | 0.31 |

<sup>a</sup>Molar mass calculated from the protein sequence after removal of the C-terminal His-tag
<sup>b</sup>Sedimentation coefficient at standard conditions (water @ 20° C) determined by combining all data from discrete component and C(S) analysis.
<sup>c</sup>Sedimentation coefficient calculated using HYDROPRO-10 from the co-ordinates of the dARC1 CA crystal structure using an atomic element radius (AER) of 2.84 Å.
<sup>d</sup>Translational diffusion coefficient for dARC1 CA determined from combined data from discrete component analysis.
<sup>e</sup>Translational diffusion coefficient calculated from the crystal structure using HYDROPRO-10, AER = 2.84 Å.
<sup>f</sup>The weight averaged molecular weight from discrete component analysis. Error is the range of three measurements
<sup>g</sup>The frictional ratio calculated from $S_{20,w}$ from discrete component analysis.
<sup>h</sup>The frictional ratio calculated from $D_{20,w}$ from discrete component analysis.
<sup>i</sup>The weight averaged molecular weight from C(S) analysis. Error is the range of three measurements
<sup>j</sup>The weight-averaged frictional ratio from the best fit C(S) distribution function.
<sup>k</sup>The range of the rms deviations observed when data were fitted using a continuous sedimentation coefficient distribution model.
<sup>l</sup>The weight averaged molecular weight from Global SE analysis. The error is the range of molecular weights observed for each multi-speed sample when fitted individually.
<sup>m</sup>The range of the rms deviations observed for each multi-speed sample when fitted individually.
<sup>n</sup>The reduced chi-squared for the global fitting of all multispeed data.

**Fig. S1.**
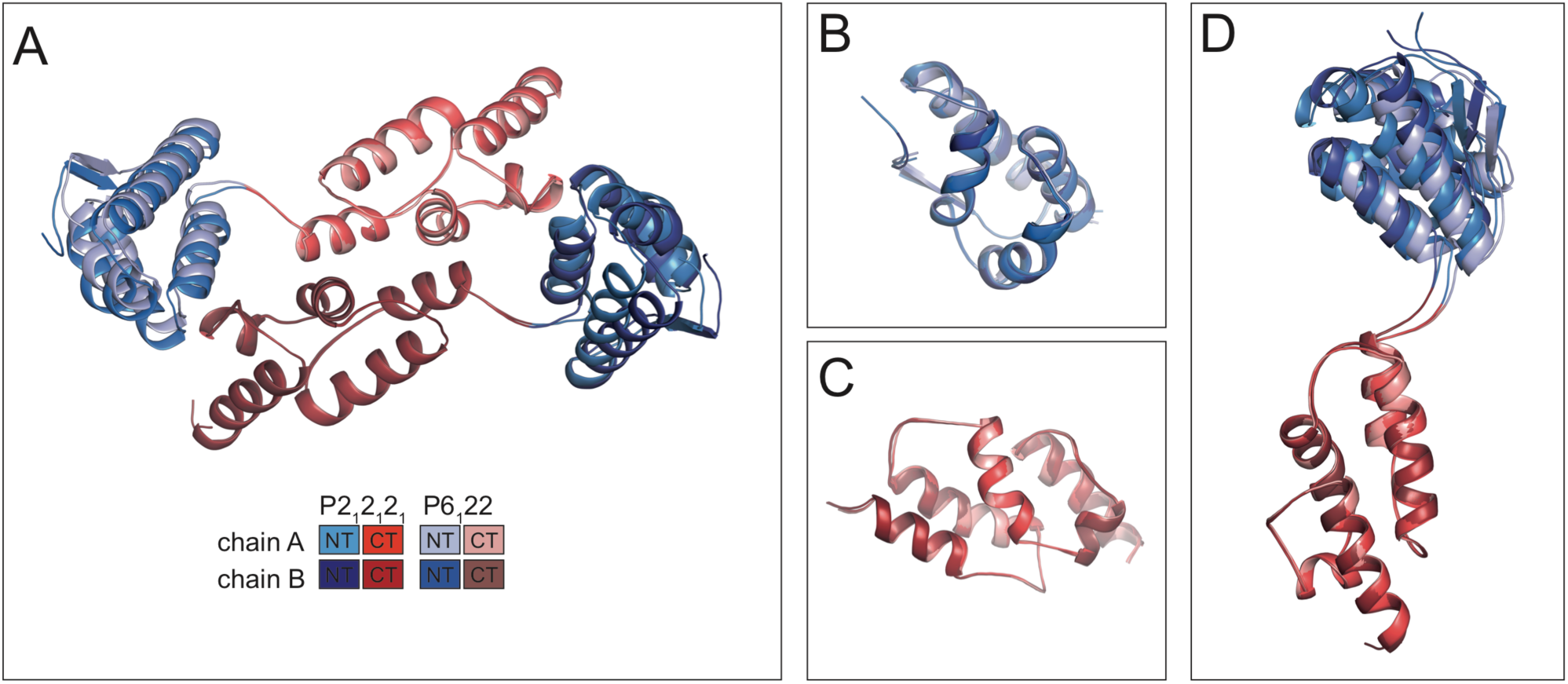
Crystal structures of the dARC1 CA. (**A**) Structural alignment of orthorhombic (P2_1_2_1_2_1_) and hexagonal (P6_1_22) crystal forms of dARC1 CA that contain an almost identical dimer. The ASUs are shown in cartoon representation, aligned using the dimeric CtDs (RMSD of 0.247 Å across 133 Cα pairs). NtDs are coloured in blue shades, CtDs in red shades, according to the legend. (**B**) Alignment of all four NtDs only, average RMSD 0.34 ± 0.09Å over 66±2 Cαs. (**C**) Alignment of all four CtDs only, average rmsd 0.32 ± 0.12 Å over 74±4 Cαs. (**D**) Alignment of all four chains through the CTD, highlighting that structural differences between chains are due to slight flexibility of the NTD-CTD linker region.

**Fig. S2.**
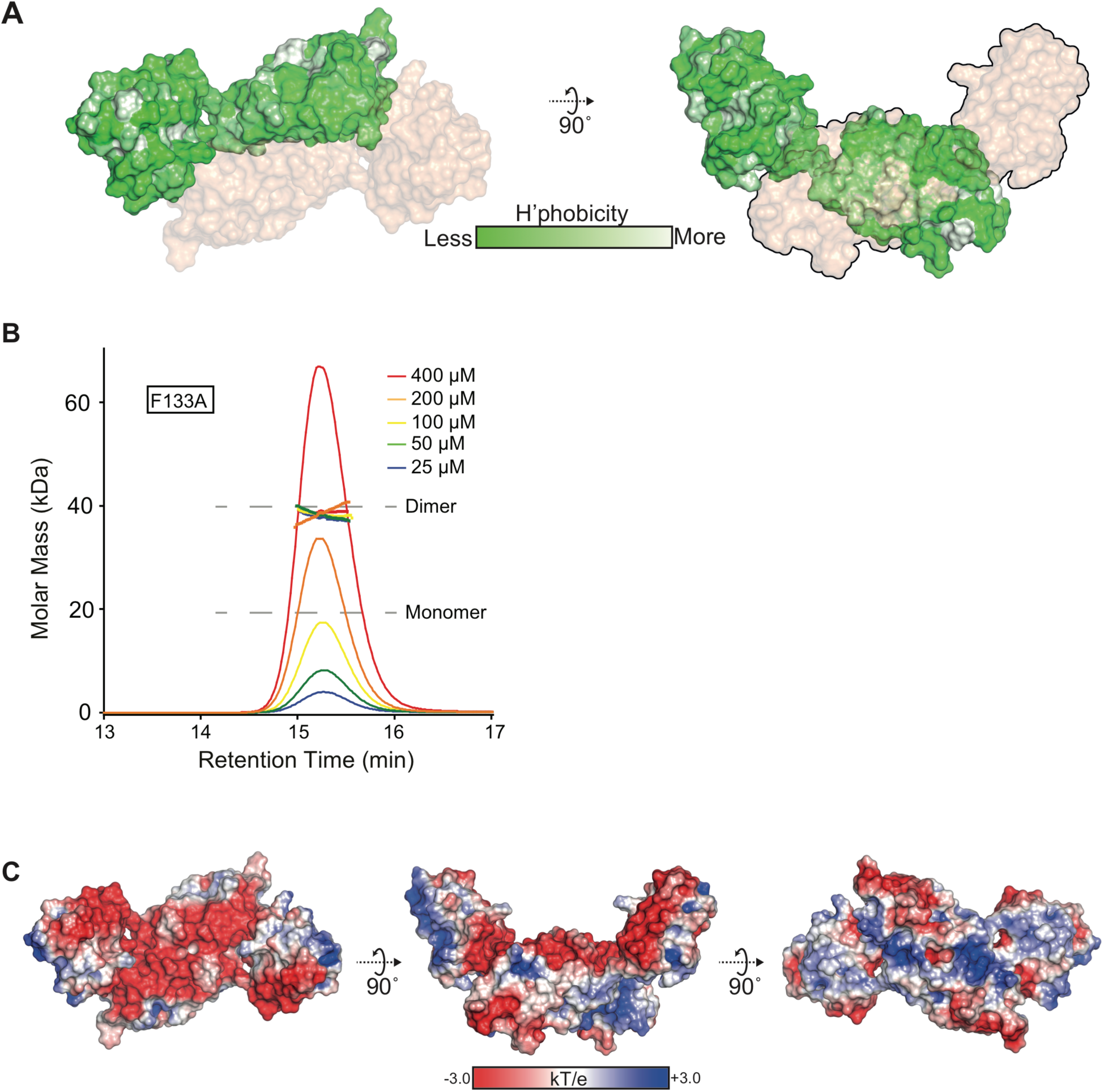
dARC1 CA dimer. (**A**) Surface representation of dARC1 CA-CtD dimer displaying the distribution of surface hydrophobicity/hydrophilicity calculated using the pymol script (https://pymolwiki.org/index.php/Color_h). Greater Hydrophilicity is represented by darker green shading, the lighter and non-coloured regions represent the most hydrophobic areas of the molecule. The orientation in the left- and right-hand panels is the same as in Fig. 1A with monomer A displaying surface hydrophobicity, and monomer B shown as a wheat surface. **(B)** SEC-MALLS analysis of dARC1 CA (F133A). The sample loading concentrations were 400 µM (8 mg/mL) (red), 200 µM (4 mg/mL) (orange), 100 µM (2 mg/mL) (yellow), 50 µM (1 mg/mL) (green) and 25 µM (0.5 mg/mL) (blue). The differential refractive index (dRI) is plotted against column retention time and the molar mass, determined at 1-second intervals throughout the elution of each peak, is plotted as points. The dARC1 CA monomer and dimer molecular mass is indicated with the grey dashed lines. (**C**) Electrostatic surface potential of the dARC1 CA dimer, calculated with APBS. The model was modified to replace SeMet with native Met residues. The view in the left-hand and centre panels is the same as in **A**. The right-hand panel is a view from the C-terminal “underside” of the dimer.

**Fig. S3.**
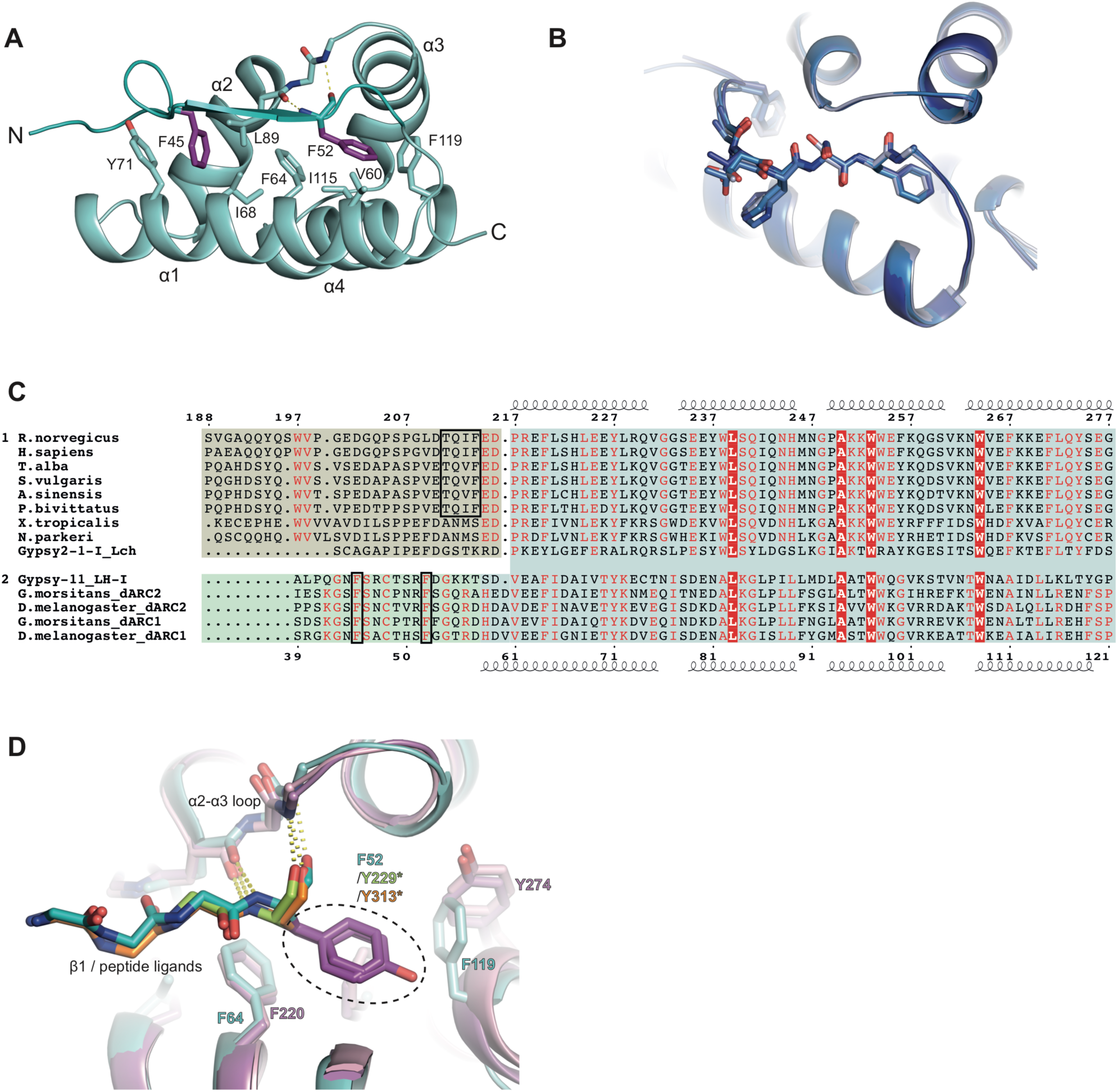
Comparison of dARC1and mam-ARC CA-NtDs. (**A**) The backbone of the dARC1 CA-NtD is shown in cartoon representation in cyan. Residues that contribute to the hydrophobic interface between the core domain, and the native N-terminal strand are shown in stick representation, coloured by atom type, with the conserved aromatic residues F45 and F52 that are buried in the interface coloured purple. The mainchain hydrogen bonding interactions between the F52 backbone amide and carbonyl with the carbonyl of L89 and the amide of Y91 are shown as dashed lines. (**B**) Overlay of the four dARC1 CA NtDs from the two crystal structures, shown in different shades of blue as in **Fig S1B**, highlighting the identical conformation of the native N-terminal strand (sticks). (**C**) Multiple sequence alignment of NtDs of ARC and dARC CA sequences. Group 1 contains tetrapod ARC (tARC) sequences and the closely related *L. ch* Gypsy2 retrotransposon. Above, rARC secondary structure, numbers according to the rARC (*R. norvegicus*) sequence). Group 2 contains dARC CA sequences and the closely related *L. h* Gypsy11 retrotransposon. Below, dARC1 secondary structure, numbers according to the dARC1 (*D. melanogaster*) sequence). Red box, white text, invariant residues shared between groups. Red text, residues conserved within a group. Blue highlight, alpha-helical core which is conserved between the two groups. Yellow highlight, native N-terminal strand which is conserved within group 1. TQIF motif is boxed in black. Green highlight, native N-terminal strand which is conserved within group 2. Conserved, strand-burying aromatic residues are boxed in black. (**D**) Close-up view of the dARC1 CA-NtD (cyan), aligned with the rARC CA NtD-CaMK2B (light pink-orange) and rARC CA NtD-TARPγ2 (dark pink-lime) structures, showing identical mechanisms of strand burial in the core domain. The native strand (dARC1) or rARC-bound peptides (TARPγ2 and CaMK2B) as well as the NtD α2-α3 loop are shown in stick representation, the equivalent strand-anchoring aromatic residue (circled) in each case is coloured purple.

**Fig. S4.**
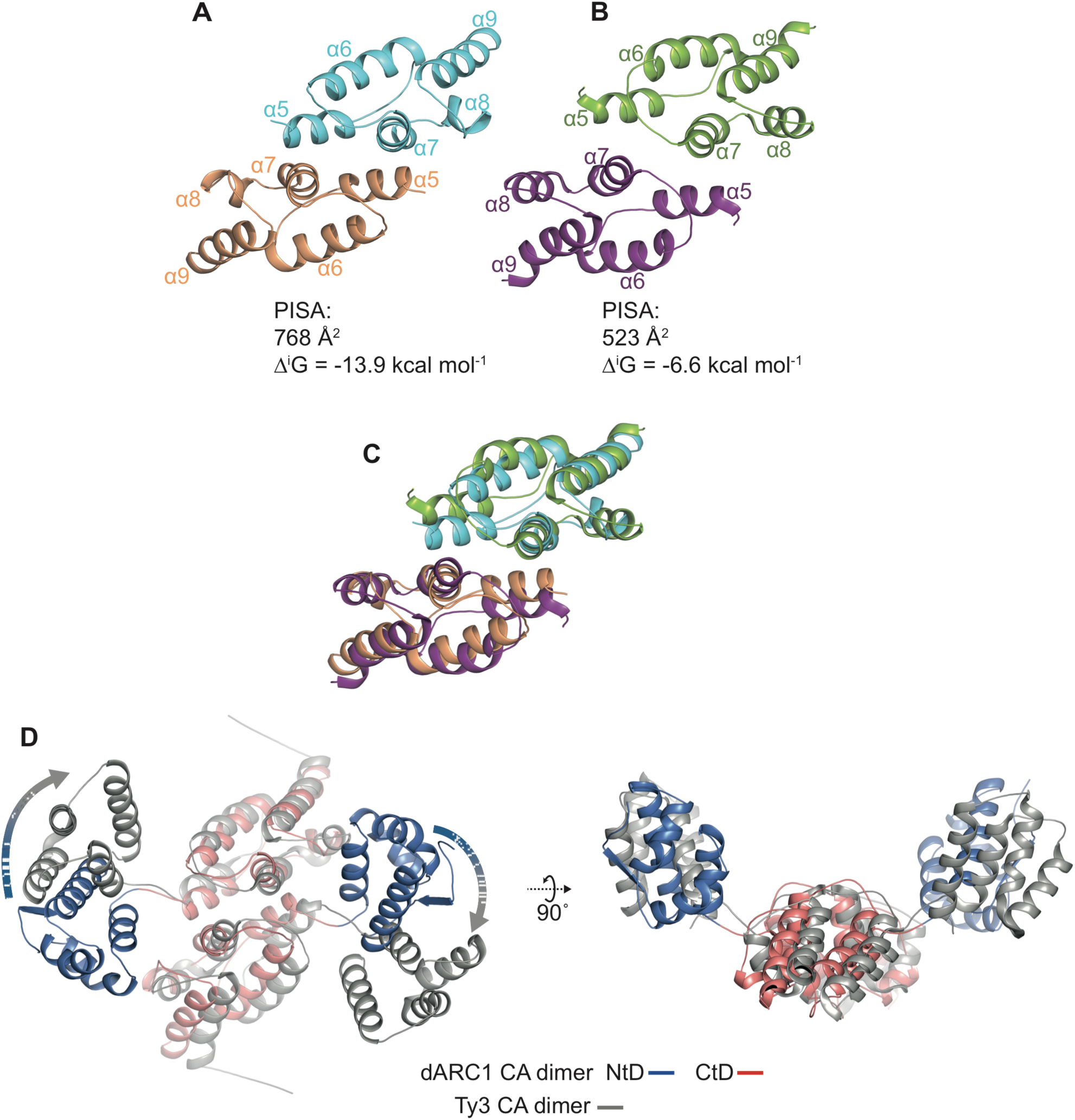
Comparison of dARC1 and Ty3 CA. **(A and B)** Cartoon representations of CA-CtD dimers. (**A**) dARC1, cyan and wheat, (**B**) Ty3 (PDB: 6r23), magenta and pale green. The structures are aligned with respect to the dARC1 dimer (RMSD = is 4.0 Å, 149 Cα). CtD helices are labelled α5 to α9 according to the dARC1 crystal structure. The buried surface area (Å^2^) and free energy of interaction (Δ^i^G) of each interface, calculated in PDBePISA is displayed below each structure. (**C**) Structural alignment of dARC1 CA-CtD and Ty3 CACtD, protein backbones are coloured as in **A and B**. (**D**) View of the dARC1 and Ty3 dimers showing the positioning of NtDs with respect to the CtD dimer. The structures are aligned on the CtD dimer as in **A**. Ty3 is coloured in grey, dARC1 CA-NtD and CA-CtD are coloured blue and red respectively. The dARC1 CA-NtDs have to undergo a slight rotational operation to match the conformation seen in the Ty3 dimer (represented by the shaded arrows). Despite this difference, both dimers have the same “glacial trough” arrangement and the dARC1 crystallographic dimer has an NtD-CtD orientation very close to that observed Ty3 icosahedral particles.

