## Supplementary material for "Structure of *D. melanogaster* ARC1 reveals a repurposed molecule with characteristics of retroviral Gag": Cottee_et_al_6S7X_PDB_report

### Full wwPDB X-ray Structure Validation Report (i)

Jul 8, 2019 – 09:17 am BST

| | | |
|---|---|---|
| PDB ID | : | 6S7X |
| Title | : | dARC1 capsid domain dimer, orthorhombic form at 1.7 Angstrom |
| Deposited on | : | 2019-07-07 |
| Resolution | : | 1.70 Å(reported) |

This is a Full wwPDB X-ray Structure Validation Report.

This report is produced by the wwPDB biocuration pipeline after annotation of the structure.

We welcome your comments at **
A user guide is available at
https://www.wwpdb.org/validation/2017/XrayValidationReportHelp
with specific help available everywhere you see the (i) symbol.

CONFIDENTIAL VALIDATION REPORT

The following versions of software and data (see references (i)) were used in the production of this report:

| | | |
|---|---|---|
| MolProbity | : | 4.02b-467 |
| Mogul | : | 1.8.0 (224370), CSD as540be (2019) |
| Xtriage (Phenix) | : | 1.13 |
| EDS | : | 2.4 |
| Percentile statistics | : | 20171227.v01 (using entries in the PDB archive December 27th 2017) |
| Refmac | : | 5.8.0158 |
| CCP4 | : | 7.0 (Gargrove) |
| Ideal geometry (proteins) | : | Engh & Huber (2001) |
| Ideal geometry (DNA, RNA) | : | Parkinson et al. (1996) |
| Validation Pipeline (wwPDB-VP) | : | 2.4 |

### 1 Overall quality at a glance (i)

The following experimental techniques were used to determine the structure:
*X-RAY DIFFRACTION*

The reported resolution of this entry is 1.70 Å.

Percentile scores (ranging between 0-100) for global validation metrics of the entry are shown in the following graphic. The table shows the number of entries on which the scores are based.

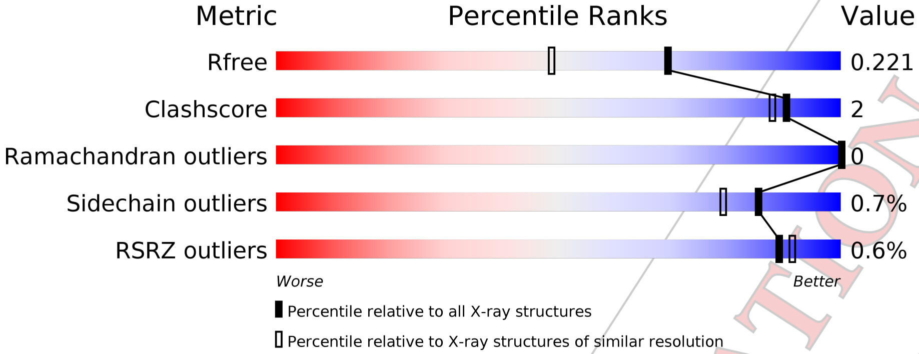


| Metric | Whole archive (#Entries) | Similar resolution (#Entries, resolution range(Å)) |
|---|---|---|
| $R_{free}$ | 111664 | 3793 (1.70-1.70) |
| Clashscore | 122126 | 4167 (1.70-1.70) |
| Ramachandran outliers | 120053 | 4100 (1.70-1.70) |
| Sidechain outliers | 120020 | 4100 (1.70-1.70) |
| RSRZ outliers | 108989 | 3718 (1.70-1.70) |

The table below summarises the geometric issues observed across the polymeric chains and their fit to the electron density. The red, orange, yellow and green segments on the lower bar indicate the fraction of residues that contain outliers for >=3, 2, 1 and 0 types of geometric quality criteria. A grey segment represents the fraction of residues that are not modelled. The numeric value for each fraction is indicated below the corresponding segment, with a dot representing fractions <=5% The upper red bar (where present) indicates the fraction of residues that have poor fit to the electron density. The numeric value is given above the bar.

| Mol | Chain | Length | Quality of chain |
|---|---|---|---|
| 1 | A | 170 | 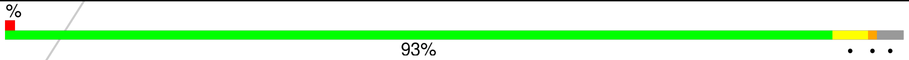 % 93% • • • |
| 1 | B | 170 | 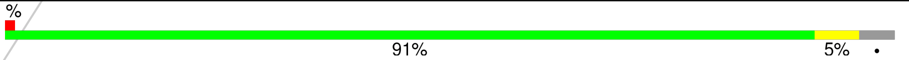 % 91% 5% • |

CONFIDENTIAL VALIDATION REPORT

### 2 Entry composition (i)

There are 4 unique types of molecules in this entry. The entry contains 5782 atoms, of which 2656 are hydrogens and 0 are deuteriums.

In the tables below, the ZeroOcc column contains the number of atoms modelled with zero occupancy, the AltConf column contains the number of residues with at least one atom in alternate conformation and the Trace column contains the number of residues modelled with at most 2 atoms.

- Molecule 1 is a protein called Activity-regulated cytoskeleton associated protein 1.

| Mol | Chain | Residues | Atoms | | | | | | | ZeroOcc | AltConf | Trace |
|---|---|---|---|---|---|---|---|---|---|---|---|---|
| | | | Total | C | H | N | O | S | Se | | | |
| 1 | A | 165 | 2725 | 877 | 1347 | 240 | 258 | 1 | 2 | 74 | 5 | 0 |
| 1 | B | 163 | 2661 | 860 | 1309 | 233 | 256 | 1 | 2 | 73 | 3 | 0 |

There are 6 discrepancies between the modelled and reference sequences:

| Chain | Residue | Modelled | Actual | Comment | Reference |
|---|---|---|---|---|---|
| A | 206 | PRO | - | expression tag | UNP Q7K1U0 |
| A | 207 | LEU | - | expression tag | UNP Q7K1U0 |
| A | 208 | GLU | - | expression tag | UNP Q7K1U0 |
| B | 206 | PRO | - | expression tag | UNP Q7K1U0 |
| B | 207 | LEU | - | expression tag | UNP Q7K1U0 |
| B | 208 | GLU | - | expression tag | UNP Q7K1U0 |

- Molecule 2 is SODIUM ION (three-letter code: NA) (formula: Na).

| Mol | Chain | Residues | Atoms | | ZeroOcc | AltConf |
|---|---|---|---|---|---|---|
| | | | Total | Na | | |
| 2 | B | 1 | 1 | 1 | 0 | 0 |
| 2 | A | 1 | 1 | 1 | 0 | 0 |

- Molecule 3 is CHLORIDE ION (three-letter code: CL) (formula: Cl).

| Mol | Chain | Residues | Atoms | | ZeroOcc | AltConf |
|---|---|---|---|---|---|---|
| | | | Total | Cl | | |
| 3 | B | 1 | 1 | 1 | 0 | 0 |
| 3 | A | 2 | 2 | 2 | 0 | 0 |

- Molecule 4 is water.

CONFIDENTIAL VALIDATION REPORT

| Mol | Chain | Residues | Atoms | ZeroOcc | AltConf |
|---|---|---|---|---|---|
| 4 | A | 201 | Total O<br>201 201 | 0 | 0 |
| 4 | B | 189 | Total O<br>190 190 | 0 | 1 |

CONFIDENTIAL VALIDATION REPORT

### 3 Residue-property plots (i)

These plots are drawn for all protein, RNA and DNA chains in the entry. The first graphic for a chain summarises the proportions of the various outlier classes displayed in the second graphic. The second graphic shows the sequence view annotated by issues in geometryand electron density. Residues are color-coded according to the number of geometric quality criteria for which they contain at least one outlier: green = 0, yellow = 1, orange = 2 and red = 3 or more. A red dot above a residue indicates a poor fit to the electron density (RSRZ > 2). Stretches of 2 or more consecutive residues without any outlier are shown as a green connector. Residues present in the sample, but not in the model, are shown in grey.

- Molecule 1: Activity-regulated cytoskeleton associated protein 1

Chain A:

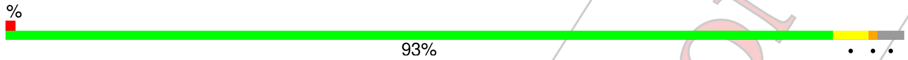


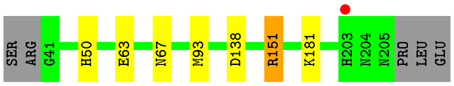


- Molecule 1: Activity-regulated cytoskeleton associated protein 1

Chain B:

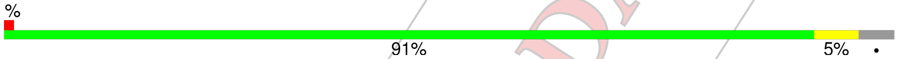


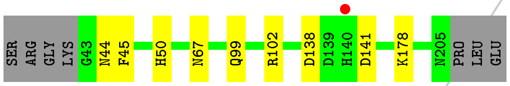


CONFIDENTIAL VALIDATION REPORT

### 4 Data and refinement statistics (i)

| Property | Value | Source |
|---|---|---|
| Space group | P 21 21 21 | Depositor |
| Cell constants<br>a, b, c, $\alpha$, $\beta$, $\gamma$ | 62.33Å 70.39Å 142.82Å<br>90.00° 90.00° 90.00° | Depositor |
| Resolution (Å) | 71.41 – 1.70<br>71.41 – 1.70 | Depositor<br>EDS |
| % Data completeness<br>(in resolution range) | 65.4 (71.41-1.70)<br>65.4 (71.41-1.70) | Depositor<br>EDS |
| $R_{merge}$ | (Not available) | Depositor |
| $R_{sym}$ | (Not available) | Depositor |
| $< I/\sigma(I) >$ [1] | 2.05 (at 1.70Å) | Xtriage |
| Refinement program | REFMAC 5.8.0238 | Depositor |
| R, $R_{free}$ | 0.181 , 0.213<br>0.192 , 0.221 | Depositor<br>DCC |
| $R_{free}$ test set | 2241 reflections (4.90%) | wwPDB-VP |
| Wilson B-factor (Å$^2$) | 20.1 | Xtriage |
| Anisotropy | 0.129 | Xtriage |
| Bulk solvent $k_{sol}$(e/Å$^3$), $B_{sol}$(Å$^2$) | 0.43 , 44.7 | EDS |
| L-test for twinning[2] | $< \|L\| > = 0.50$, $< L^2 > = 0.33$ | Xtriage |
| Estimated twinning fraction | No twinning to report. | Xtriage |
| $F_o$,$F_c$ correlation | 0.95 | EDS |
| Total number of atoms | 5782 | wwPDB-VP |
| Average B, all atoms (Å$^2$) | 24.0 | wwPDB-VP |

Xtriage's analysis on translational NCS is as follows: *The largest off-origin peak in the Patterson function is 5.44% of the height of the origin peak. No significant pseudotranslation is detected.*

[1] Intensities estimated from amplitudes.

[2] Theoretical values of $< |L| >$, $< L^2 >$ for acentric reflections are 0.5, 0.333 respectively for untwinned datasets, and 0.375, 0.2 for perfectly twinned datasets.

CONFIDENTIAL VALIDATION REPORT

### 5 Model quality (i)

#### 5.1 Standard geometry (i)

Bond lengths and bond angles in the following residue types are not validated in this section: NA, CL

The Z score for a bond length (or angle) is the number of standard deviations the observed value is removed from the expected value. A bond length (or angle) with $|Z| > 5$ is considered an outlier worth inspection. RMSZ is the root-mean-square of all Z scores of the bond lengths (or angles).

| Mol | Chain | Bond lengths | | Bond angles | |
|---|---|---|---|---|---|
| | | RMSZ | $\#\|Z\| >5$ | RMSZ | $\#\|Z\| >5$ |
| 1 | A | 0.71 | 0/1421 | 0.87 | 2/1913 (0.1%) |
| 1 | B | 0.69 | 0/1392 | 0.80 | 0/1876 |
| All | All | 0.70 | 0/2813 | 0.83 | 2/3789 (0.1%) |

There are no bond length outliers.

All (2) bond angle outliers are listed below:

| Mol | Chain | Res | Type | Atoms | Z | Observed($^o$) | Ideal($^o$) |
|---|---|---|---|---|---|---|---|
| 1 | A | 93 | MSE | CG-SE-CE | -9.37 | 78.29 | 98.90 |
| 1 | A | 181 | LYS | CB-CA-C | -5.15 | 100.10 | 110.40 |

There are no chirality outliers.

There are no planarity outliers.

#### 5.2 Too-close contacts (i)

In the following table, the Non-H and H(model) columns list the number of non-hydrogen atoms and hydrogen atoms in the chain respectively. The H(added) column lists the number of hydrogen atoms added and optimized by MolProbity. The Clashes column lists the number of clashes within the asymmetric unit, whereas Symm-Clashes lists symmetry related clashes.

| Mol | Chain | Non-H | H(model) | H(added) | Clashes | Symm-Clashes |
|---|---|---|---|---|---|---|
| 1 | A | 1378 | 1347 | 1340 | 6 | 0 |
| 1 | B | 1352 | 1309 | 1302 | 4 | 0 |
| 2 | A | 1 | 0 | 0 | 0 | 0 |
| 2 | B | 1 | 0 | 0 | 0 | 0 |
| 3 | A | 2 | 0 | 0 | 0 | 0 |
| 3 | B | 1 | 0 | 0 | 0 | 0 |
| 4 | A | 201 | 0 | 0 | 4 | 1 |

*Continued on next page...*

CONFIDENTIAL VALIDATION REPORT

*Continued from previous page...*

| Mol | Chain | Non-H | H(model) | H(added) | Clashes | Symm-Clashes |
|---|---|---|---|---|---|---|
| 4 | B | 190 | 0 | 0 | 0 | 0 |
| All | All | 3126 | 2656 | 2642 | 10 | 1 |

The all-atom clashscore is defined as the number of clashes found per 1000 atoms (including hydrogen atoms). The all-atom clashscore for this structure is 2.

All (10) close contacts within the same asymmetric unit are listed below, sorted by their clash magnitude.

| Atom-1 | Atom-2 | Interatomic distance (Å) | Clash overlap (Å) |
|---|---|---|---|
| 1:A:151[A]:ARG:HD2 | 4:A:476:HOH:O | 1.82 | 0.79 |
| 1:A:151[A]:ARG:HH21 | 1:A:151[A]:ARG:HB3 | 1.59 | 0.66 |
| 1:A:63:GLU:OE2 | 4:A:401:HOH:O | 2.14 | 0.66 |
| 1:A:151[A]:ARG:CD | 4:A:476:HOH:O | 2.43 | 0.66 |
| 1:A:50:HIS:CD2 | 1:A:67:ASN:HB3 | 2.33 | 0.64 |
| 1:B:99[A]:GLN:OE1 | 1:B:102:ARG:NH2 | 2.35 | 0.59 |
| 1:B:44:ASN:O | 1:B:45:PHE:HB2 | 2.11 | 0.50 |
| 1:B:138:ASP:HB2 | 1:B:141:ASP:HB2 | 1.98 | 0.46 |
| 1:A:138:ASP:HB3 | 4:A:517:HOH:O | 2.17 | 0.44 |
| 1:B:50:HIS:CD2 | 1:B:67:ASN:HB3 | 2.55 | 0.42 |

All (1) symmetry-related close contacts are listed below. The label for Atom-2 includes the symmetry operator and encoded unit-cell translations to be applied.

| Atom-1 | Atom-2 | Interatomic distance (Å) | Clash overlap (Å) |
|---|---|---|---|
| 4:A:502:HOH:O | 4:A:513:HOH:O[4_556] | 2.09 | 0.11 |

#### 5.3 Torsion angles (i)

##### 5.3.1 Protein backbone (i)

In the following table, the Percentiles column shows the percent Ramachandran outliers of the chain as a percentile score with respect to all X-ray entries followed by that with respect to entries of similar resolution.

The Analysed column shows the number of residues for which the backbone conformation was analysed, and the total number of residues.

| Mol | Chain | Analysed | Favoured | Allowed | Outliers | Percentiles | |
|---|---|---|---|---|---|---|---|
| 1 | A | 168/170 (99%) | 168 (100%) | 0 | 0 | 100 | 100 |

*Continued on next page...*

CONFIDENTIAL VALIDATION REPORT

*Continued from previous page...*

| Mol | Chain | Analysed | Favoured | Allowed | Outliers | Percentiles | |
|---|---|---|---|---|---|---|---|
| 1 | B | 164/170 (96%) | 162 (99%) | 2 (1%) | 0 | 100 | 100 |
| All | All | 332/340 (98%) | 330 (99%) | 2 (1%) | 0 | 100 | 100 |

There are no Ramachandran outliers to report.

##### 5.3.2 Protein sidechains (i)

In the following table, the Percentiles column shows the percent sidechain outliers of the chain as a percentile score with respect to all X-ray entries followed by that with respect to entries of similar resolution.

The Analysed column shows the number of residues for which the sidechain conformation was analysed, and the total number of residues.

| Mol | Chain | Analysed | Rotameric | Outliers | Percentiles | |
|---|---|---|---|---|---|---|
| 1 | A | 149/147 (101%) | 147 (99%) | 2 (1%) | 71 | 58 |
| 1 | B | 146/147 (99%) | 145 (99%) | 1 (1%) | 85 | 79 |
| All | All | 295/294 (100%) | 292 (99%) | 3 (1%) | 85 | 69 |

All (3) residues with a non-rotameric sidechain are listed below:

| Mol | Chain | Res | Type |
|---|---|---|---|
| 1 | A | 151[A] | ARG |
| 1 | A | 151[B] | ARG |
| 1 | B | 178 | LYS |

Some sidechains can be flipped to improve hydrogen bonding and reduce clashes. All (1) such sidechains are listed below:

| Mol | Chain | Res | Type |
|---|---|---|---|
| 1 | A | 50 | HIS |

##### 5.3.3 RNA (i)

There are no RNA molecules in this entry.

#### 5.4 Non-standard residues in protein, DNA, RNA chains (i)

There are no non-standard protein/DNA/RNA residues in this entry.

CONFIDENTIAL VALIDATION REPORT

#### 5.5 Carbohydrates (i)

There are no carbohydrates in this entry.

#### 5.6 Ligand geometry (i)

Of 5 ligands modelled in this entry, 5 are monoatomic - leaving 0 for Mogul analysis.

There are no bond length outliers.

There are no bond angle outliers.

There are no chirality outliers.

There are no torsion outliers.

There are no ring outliers.

No monomer is involved in short contacts.

#### 5.7 Other polymers (i)

There are no such residues in this entry.

#### 5.8 Polymer linkage issues (i)

There are no chain breaks in this entry.

CONFIDENTIAL VALIDATION REPORT

### 6 Fit of model and data (i)

#### 6.1 Protein, DNA and RNA chains (i)

In the following table, the column labelled '#RSRZ> 2' contains the number (and percentage) of RSRZ outliers, followed by percent RSRZ outliers for the chain as percentile scores relative to all X-ray entries and entries of similar resolution. The OWAB column contains the minimum, median, $95^{th}$ percentile and maximum values of the occupancy-weighted average B-factor per residue. The column labelled 'Q< 0.9' lists the number of (and percentage) of residues with an average occupancy less than 0.9.

| Mol | Chain | Analysed | <RSRZ> | #RSRZ>2 | | | OWAB(Å$^2$) | Q<0.9 |
|---|---|---|---|---|---|---|---|---|
| 1 | A | 163/170 (95%) | -0.07 | 1 (0%) | 89 | 91 | 12, 20, 36, 59 | 0 |
| 1 | B | 161/170 (94%) | -0.06 | 1 (0%) | 89 | 91 | 12, 22, 43, 70 | 0 |
| All | All | 324/340 (95%) | -0.07 | 2 (0%) | 89 | 91 | 12, 21, 41, 70 | 0 |

All (2) RSRZ outliers are listed below:

| Mol | Chain | Res | Type | RSRZ |
|---|---|---|---|---|
| 1 | A | 203 | HIS | 2.4 |
| 1 | B | 140 | HIS | 2.4 |

#### 6.2 Non-standard residues in protein, DNA, RNA chains (i)

There are no non-standard protein/DNA/RNA residues in this entry.

#### 6.3 Carbohydrates (i)

There are no carbohydrates in this entry.

#### 6.4 Ligands (i)

In the following table, the Atoms column lists the number of modelled atoms in the group and the number defined in the chemical component dictionary. The B-factors column lists the minimum, median, $95^{th}$ percentile and maximum values of B factors of atoms in the group. The column labelled 'Q< 0.9' lists the number of atoms with occupancy less than 0.9.

| Mol | Type | Chain | Res | Atoms | RSCC | RSR | B-factors(Å$^2$) | Q<0.9 |
|---|---|---|---|---|---|---|---|---|
| 3 | CL | A | 303 | 1/1 | 0.98 | 0.06 | 28,28,28,28 | 0 |
| 2 | NA | A | 301 | 1/1 | 0.99 | 0.12 | 39,39,39,39 | 0 |
| 3 | CL | B | 302 | 1/1 | 0.99 | 0.10 | 37,37,37,37 | 0 |

*Continued on next page...*

CONFIDENTIAL VALIDATION REPORT

*Continued from previous page...*

| Mol | Type | Chain | Res | Atoms | RSCC | RSR | B-factors(Å$^2$) | Q<0.9 |
|---|---|---|---|---|---|---|---|---|
| 2 | NA | B | 301 | 1/1 | 0.99 | 0.07 | 33,33,33,33 | 0 |
| 3 | CL | A | 302 | 1/1 | 1.00 | 0.06 | 27,27,27,27 | 0 |

#### 6.5 Other polymers (i)

There are no such residues in this entry.

CONFIDENTIAL VALIDATION REPORT
